# Insertions in the SARS-CoV-2 Spike N-Terminal Domain May Aid COVID-19 Transmission

**DOI:** 10.1101/2021.12.06.471394

**Authors:** Su Datt Lam, Vaishali P Waman, Christine Orengo, Jonathan Lees

**Author notes:** **Correspondence:** SDL; CO; JL.

## Abstract

Coronavirus disease 2019 (COVID-19) caused by SARS-CoV-2 is an ongoing pandemic that causes significant health/socioeconomic burden. Variants of concern (VOCs) have emerged which may affect transmissibility, disease severity and re-infection risk. Most studies focus on the receptor-binding domain (RBD) of the Spike protein. However, some studies suggest that the Spike N-terminal domain (NTD) may have a role in facilitating virus entry via sialic-acid receptor binding. Furthermore, most VOCs include novel NTD variants. Recent analyses demonstrated that NTD insertions in VOCs tend to lie close to loop regions likely to be involved in binding sialic acids. We extended the structural characterisation of these putative sugar binding pockets and explored whether variants could enhance the binding to sialic acids and therefore to the host membrane, thereby contributing to increased transmissibility. We found that recent NTD insertions in VOCs (i.e., Gamma, Delta and Omicron variants) and emerging variants of interest (VOIs) (i.e., Iota, Lambda, Theta variants) frequently lie close to known and putative sugar-binding pockets. For some variants, including the recent Omicron VOC, we find increases in predicted sialic acid binding energy, compared to the original SARS-CoV-2, which may contribute to increased transmission. We examined the similarity of NTD across a range of related Betacoronaviruses to determine whether the putative sugar-binding pockets are sufficiently similar to be exploited in drug design. Despite global sequence and structure similarity, most sialic-acid binding pockets of NTD vary across related coronaviruses. Typically, SARS-CoV-2 possesses additional loops in these pockets that increase contact with polysaccharides. Our work suggests ongoing evolutionary tuning of the sugar-binding pockets in the virus. Whilst three of the pockets are too structurally variable to be amenable to pan Betacoronavirus drug design, we detected a fourth pocket that is highly structurally conserved and could therefore be investigated in pursuit of a generic drug. Our structure-based analyses help rationalise the effects of VOCs and provide hypotheses for experiments. For example, the Omicron variant, which has increased binding to sialic acids in pocket 3, has a rather unique insertion near pocket 3. Our work suggests a strong need for experimental monitoring of VOC changes in NTD.

## 1 Introduction

Coronavirus disease 2019 (COVID-19), first detected in December 2019, is an ongoing pandemic situation and a cause of significant health and socioeconomic burden globally. COVID-19 is caused by a novel coronavirus namely SARS-CoV-2, which has led to the loss of the lives of around 4.2 million people worldwide (World Health Organization, 2021).

Vaccines are being successfully administered, however the emergence of new variants of concern (VOCs) raises key questions about their phenotypic impact on transmissibility, disease severity, risk of re-infection and impact on diagnostics (World Health Organization, 2021). So far, five VOCs have been detected namely: Alpha variant (reported in 197 countries), Beta (147 countries), Gamma (105 countries), Delta (in 201 countries) and Omicron variant (in 19 countries) (World Health Organization, 2021). All VOCs are reported to carry novel mutations particularly in the domains of their spike protein, along with a few additional mutations in other parts of the virus genome (http://sars2.cvr.gla.ac.uk/cog-uk/;https://www.who.int/en/activities/tracking-SARS-CoV-2-variants/).

The Spike protein (S) is the key antigen and plays a major role in virus host cell attachment and entry and fusion with the host cell membrane. It is the primary target for neutralising antibodies during infection and the majority of vaccines. It is a homo-trimeric protein and in SARS-CoV-2, it is made up of two subunits, S1 and S2 (Xu et al., 2021). The S1 subunit comprises two domains: the NTD (N-terminal domain) and the RBD (receptor binding domain). Structural studies have revealed mechanisms by which the RBD of the SARS-CoV-2 spike protein interacts with human ACE2 protein to gain entry into the host cell (Lan et al., 2020; Shang et al., 2020).

A recent study by Dicken and co-workers (2021) showed that the N-terminal domain (NTD) of the spike protein might also have a role in facilitating efficient virus entry. Their experimental study of the Alpha variant investigated how amino acid deletions in NTD resulted in more efficient entry into human cells and infection, as compared to the reference strain (Wuhan-Hu-1). Furthermore, the NTD domain is observed to be a sequence diversity hotspot. The comprehensive sequence-based analyses of the Sarbecovirus subgenus of the Betacoronaviruses (BCoVs) by Dicken et al. (2021) showed five distinct regions insertions and deletions (indels) in the NTD domains of different BCoVs. These were found to be associated with flexible loops in the NTD structure. Intriguingly, indel regions 1, 3 and 5 correspond to an NTD antigenic supersite (McCallum et al., 2021) (see Figure 4B), that is a target of all known NTD-specific neutralising antibodies, thus supporting the role of NTD in eliciting protective immunity. McCallum et al. (2021) recently revealed that most of the VOCs harbour novel mutations in this NTD antigenic supersite.

The NTD domain of the Spike protein has been proposed to bind to C-type lectin receptors such as L-SIGN/CD-SIGN (Amraei et al., 2020). Furthermore, the role of NTD in binding to sialic acid-containing glycoproteins and gangliosides has been reported in several coronaviruses and other viruses (Fantini et al., 2020; Seyran et al., 2021). There is also some evidence from experimental studies of other BCoVs that changes in NTD may improve affinity for sugars on the host cell surface and therefore increase the affinity of the virus for the specific host cell (Hulswit et al., 2019; Qing et al., 2020). Sialic acids comprise a nine-carbon backbone and are located at the terminal end of glycans. Several in silico structural studies have searched for and analysed pockets in NTD. Three pockets have been found, to date, two of which have been shown to bind sialic acid (Awasthi et al., 2020; Fantini et al., 2020; Robson, 2020; Baker et al., 2021; Bò et al., 2021; Di Gaetano et al., 2021; Milanetti et al., 2021).

The N-terminal domain of SARS-CoV-2 Spike protein is a beta sandwich protein belonging to the “Spike glycoprotein, N-terminal domain” superfamily (ID: 2.60.120.960) in the CATH structural classification (Sillitoe et al., 2021). The sugar-binding protein human Galectin is also found within the same fold group (2.60.120) and the two domains are structurally similar. Studies have shown that the NTD of HKU-23 (a BCoV) is structurally related to the Galectin fold (e.g., Galectin-3), possibly co-opted by the virus from the host (Li, 2016). More recently, Cheng et al. showed that BCoVs (BovineCoV-NTD, PHEV-NTD, HCoV-OC43-NTD, HCoV-HKU23-NTD, HKU1-NTD, MHV-NTD, SARS-CoV-NTD and MERS-NTD) bind to sugars such as sialic acid-containing glycoproteins and gangliosides in an equivalent binding pocket to Galectin-3 (referred to as ‘Pocket 1’) (Cheng et al., 2019).

Interestingly, Behloul and co-workers found a GTNGTKR motif, a known sugar binding motif (located at residues 72-78 in SARS-CoV-2 Spike protein) within a different pocket on the protein (referred to as ‘Pocket 2’) suggesting that this region may also bind sugars (Behloul et al., 2020). Awasthi et al (2020) performed a comparative structural analysis of pocket 2 in the NTD of SARS-CoV-2 and SARS-CoV, using molecular dynamics and docking approaches. Their analyses highlighted the flexible nature of the loop (L244-G261, part of pocket 2) allowing pocket 2 of SARS-CoV-2:NTD to bind sialosides more strongly than that of SARS-CoV (Awasthi et al., 2020). However, human Galectin-3 does not bind sugar in this pocket 2 region, it solely utilises pocket 1 (Johannes et al., 2018). Pocket 2 is detected in other coronaviruses including BovineCoV-NTD, PHEV-NTD, HCoV-OC43-NTD, HCoV-HKU23-NTD, HKU1-NTD, MHV-NTD but not well defined in SARS-CoV-NTD nor MERS-NTD (Cheng et al., 2019). Furthermore, the structure of HCoV-OC43 has been determined with a sialic acid (i.e., 9-O-acetyl-sialic acid) bound in Pocket 2 (Tortorici et al., 2019). More recently, a third druggable pocket was detected in NTD domain of SARS-CoV-2, but there is no indication to date on whether it can bind sugars (Di Gaetano et al., 2021).

Our work builds on these previous studies and further explores the role of NTD in sialic acid binding. We explore the characteristics of all putative sialic binding pockets in NTD and examine the effects of variants (residue insertions/deletions) in these pockets on sialic acid binding. In particular we use docking studies to characterise the pockets and observe that pockets 2 and 3 bind sialic acid more strongly than pocket 1. We also detect a fourth pocket (pocket 4) that can bind sialic acid but not as strongly as pockets 2 and 3.

Since Awasthi et al. (2020) showed that insertions in SARS-CoV-2 relative to SARS-CoV enhanced sialic binding, we structurally compared all 4 of the pockets across related coronaviruses to determine whether (and if so how) these pockets had evolved to enhance binding to polysaccharides. Our results support the work of Awasthi et al, showing that SARS-CoV-2 possesses additional loops in pocket 2 that extend the pocket to increase contact with polysaccharides. We also found extensions in pocket 3. We then determined whether variants, found in VOCs and VOIs of SARS-CoV-2, including insertions/deletions in the loops defining the NTD sugar binding pockets could result in enhanced binding of the spike protein to sialic acid (and therefore to the host membrane).

Finally, we assessed which of the pockets were druggable and sufficiently structurally conserved across related coronaviruses for their structural features to be exploited in the design of generic drugs against these BCoV viruses. Whilst there are some therapeutic strategies targeted at pocket 1 in SARS-CoV (Fantini et al., 2020), this pocket is too variable across BCoVs. Pocket 2 and 3 are even more structurally variable and again not amenable to generic drug design. However, the fourth druggable pocket, pocket 4, appears to be highly structurally conserved and could therefore be further investigated in pursuit of a generic (pan BCoVs) drug. Sialic acid targeting, or mimicking drugs could serve as good candidates for antiviral strategies (Heida et al., 2021).

## 2 Materials and Methods

### 2.1 Sequence data

To explore the sequence and structure conservation of BCoVs, we selected a set of representative sequences, expected to be closely related to SARS-CoV-2 and also strains that have caused human disease (Zhao et al., 2020). We obtained 33 nucleotide sequences of the Sarbecovirus subgenus of the BCoV and SARS-related coronaviruses from NCBI (Sayers et al., 2020) and GISAID (Elbe and Buckland-Merrett, 2017; Shu and McCauley, 2017) databases (See Supplementary Table 1). We extracted the Spike protein sequences by scanning the sequence of the SARS-CoV-2 Spike protein (YP_009724390.1) against the nucleotide database of BCoV sequences using NCBI BLAST v2.6 tblastn (Altschul et al., 1990). For each of the Spike protein sequences extracted, we then extracted the N-terminal domain (residues 1-303).

We also obtained BCoV NTD domain sequences for these proteins from the CATH “Spike glycoprotein, N-terminal domain” domain superfamily (2.60.120.960) (Sillitoe et al., 2021).

### 2.2 Structure data

We obtained experimental structures for the NTD-domain of SARS-CoV (PDB ID: 6ACC), Pangolin-CoV-GX (PDB ID: 7CN8), Pangolin-CoV-GD (PDB ID: 7BBH), Bat-Cov-RaTG13 (PDB ID: 7CN4), SARS-COV-2 (PDB ID: 7C2L and PDB ID: 6ZGE), HCoV-OC43 (PDB ID: 6NZK) and Human Galectin-3 (PDB ID: 1A3K) from the Protein Data Bank (PDB) (Berman et al., 2000).

Structural models of other BCoV NTDs were built using an in-house FunMOD modelling pipeline (Lam et al., 2017; Sillitoe et al., 2020), provided they had more than 40% sequence identity to the template (see Supplementary Table 2 for details). Otherwise, models were built using AlphaFold2 (Jumper et al., 2021, ‘AlphaFold2_mmseqs2’ notebook available from https://github.com/sokrypton/ColabFold, Mirdita et al., 2021) (see Supplementary Table 3 for details). FunMOD generated query–template alignments using HH-suite version 3 (Steinegger et al., 2019), which were then used as input to the MODELLER v.9.23 program (Webb and Sali, 2016). We used the ‘very_slow’ schedule for model refinement. Ten models were generated for each query and we then selected the model with the lowest normalised DOPE score (nDOPE) (Shen and Sali, 2006), which reflects the quality of the model. Positive scores are likely to be poor models, while scores lower than −1 are likely to be native-like.

### 2.3 Multiple sequence alignment and phylogenetic tree

To build a Maximum likelihood tree, we aligned all NTD amino acid sequences using CLUSTAL-OMEGA (Madeira et al., 2019). A phylogenetic tree was inferred according to the Maximum Likelihood method. Genetic distance was computed using the Whelan and Goldman model (Whelan and Goldman, 2001) and gamma-distributed rate variation among sites (WAG + G). The evolutionary history was inferred using the Maximum Likelihood method and the analyses were conducted using MEGA X (Kumar et al., 2018).

We used ESPript3 webserver (http://espript.ibcp.fr/ESPript/ESPript/)(Robert and Gouet, 2014) to produce structure-based multiple sequence alignment. The secondary structure elements were defined based on the SARS-CoV-2 NTD structure (PDB ID 6ZGE).

The ScoreCons method (Valdar, 2002) was used to calculate the sequence conservation of residues at a particular position. ScoreCons reports a score between 0-1 which is robust for multiple sequence alignments with high information content (DOPS score >70). A ScoreCons value above 0.7 suggests a highly conserved position.

### 2.4 Structure Comparison and Structural Analysis

Protein structures were compared using our in-house SSAP algorithm (Taylor and Orengo, 1989). The SSAP score ranges from 0 to 100. Structures with a SSAP score above 80 are considered to be highly similar. Protein structures were rendered using PyMOL (Schrödinger and DeLano, 2020) and UCSF Chimera (Pettersen et al., 2004).

### 2.5 Druggable pocket prediction and Molecular docking

We used CavityPlus (Xu et al., 2018) to predict druggable pockets using chain A of SARS-CoV-2 structure (PDB ID: 7C2L). The CavityPlus DrugScore denotes the druggability of a particular pocket with more positive scores indicating more druggable sites.

Molecular docking was done using the HADDOCK 2.4 webserver (https://wenmr.science.uu.nl/haddock2.4/) (Honorato et al., 2021). HADDOCK was one of the top-performing protein-ligand binding methods in the D3R Grand Challenge 2 and 3 (Kurkcuoglu et al., 2018; Gaieb et al., 2019). The sialic acid used was 9-O-acetylated sialic acid (PubChem ID 71312953). This ligand is the sialic acid bound to the experimental structure of HCoV-OC43 (PDB ID 6NZK, Tortorici et al., 2019). The HADDOCK score is a linear combination of van der Waals, electrostatics, and desolvation energy terms. A lower HADDOCK score signifies stronger binding. The binding affinity of sialic acid-NTD was predicted using HADDOCK’s PRODIGY-LIGAND server (https://wenmr.science.uu.nl/prodigy/) (Vangone et al., 2019).

We used the LIGPLOT program of PDBsum (Laskowski and Swindells, 2011) to examine residue interactions between sialic acid and the NTDs.

## 3 Results

### 3.1 Phylogenetic studies of coronavirus NTD domains

We performed phylogenetic studies on the NTD domain to explore the relationships between the different BCoVs. We obtained sequences of the NTD domain for 26 BCoVs from the NCBI GenBank database and GISAID database.

BCoVs have been classified into 5 lineages: Embecovirus (lineage A), Sarbecovirus (lineage B), Merbecovirus (lineage C), Nobecovirus (lineage D) and Hibecovirus (Woo et al., 2010; Zhao et al., 2020). SARS-CoV-2 and SARS-related coronaviruses belong to the Sarbecovirus lineage (lineage B). HCoV-OC43 belongs to Embecovirus (lineage A), while MERS-CoV was classified into Merbecovirus (lineage C). So far, there have been no reports of human-infecting BCoVs from other lineages.

Figure 1 demonstrates that the phylogenetic tree of SARS-CoV-2 based on the NTD domain, is similar to those constructed from other parts of the Spike protein i.e. SARS-CoV-2, BatCOV-RaTG13, PangolinCoV and BatCoV-ZXC21 and BatCoV-ZC45 fall into the same clade in the tree. In contrast, SARS-CoV, Civet-SARSr-CoV and BatCoV-RmYN02 were found to belong to another clade. HCoV-OC43, which is thought to have emerged in the 1950s (Vijgen et al., 2005; Lau et al., 2011) and MERS-CoV are more distant from the others. See Supplementary Figure 1 for a bootstrap tree.

**Figure 1.**
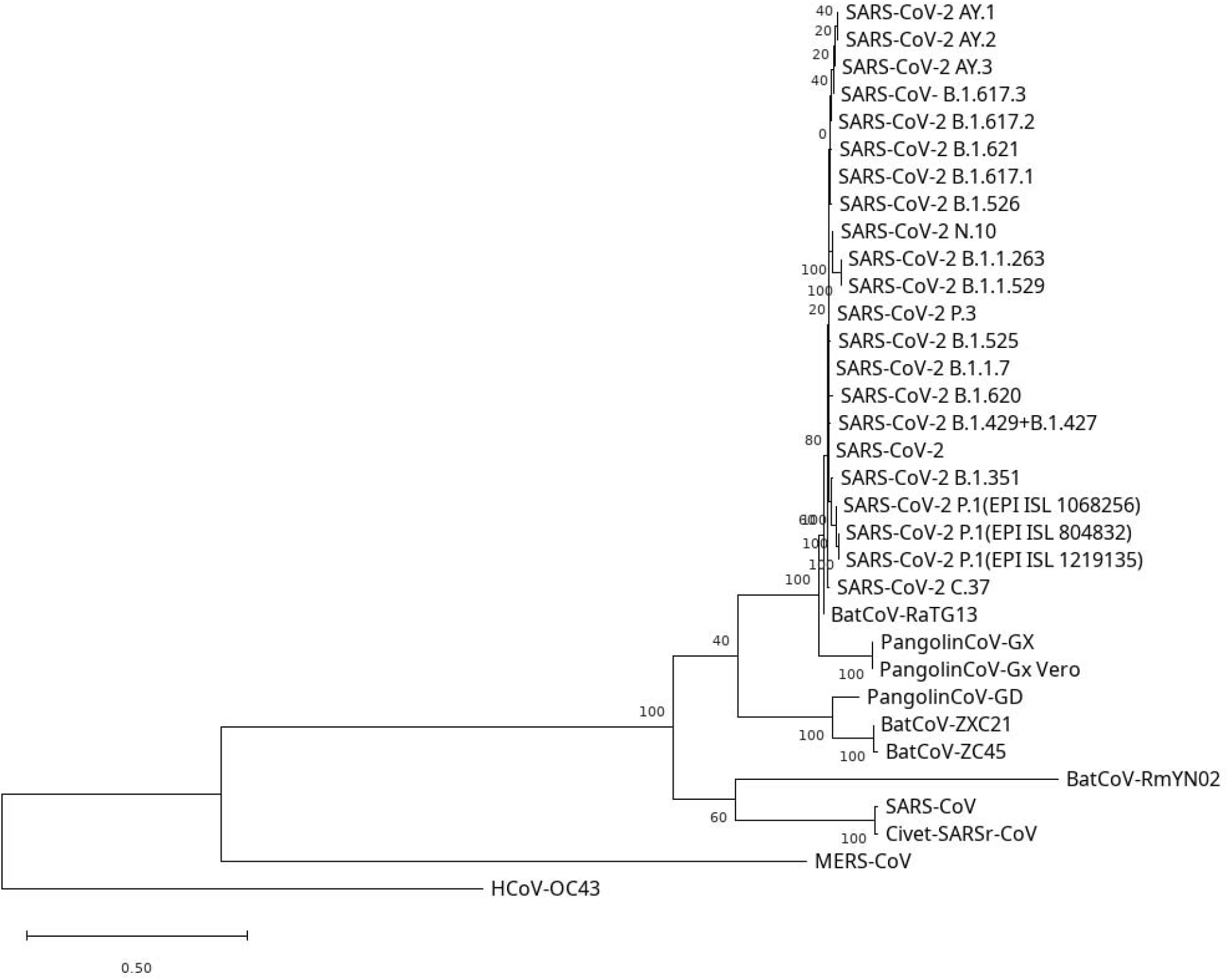
Phylogenetic tree of BCoV NTD amino acid sequences. The phylogenetic tree was inferred according to the Maximum Likelihood method. Genetic distance was computed using the Whelan and Goldman model and gamma-distributed rate variation among sites (WAG + G).

### 3.2 Analyses of structural similarity between the coronavirus NTD domains

The structures of the Spike proteins from the Wuhan strain of SARS-CoV-2 and several other strains (i.e. SARS-CoV, PangolinCoV-GX, PangolinCoV-GD, BatCov-RaTG13) have been determined by experimental methods (Song et al., 2018; Wrapp et al., 2020; Wrobel et al., 2021; Zhang et al., 2021). The NTD domain interacts with a number of other domains within the Spike protein (subdomains SD1 and SD2, which serve as a hinge for RBD up-movement (Berger and Schaffitzel, 2020; Peters et al., 2020)), however a significant proportion of the domain is surface accessible and able to interact with other compounds and proteins on the host cell surface (see Figure 2).

**Figure 2.**
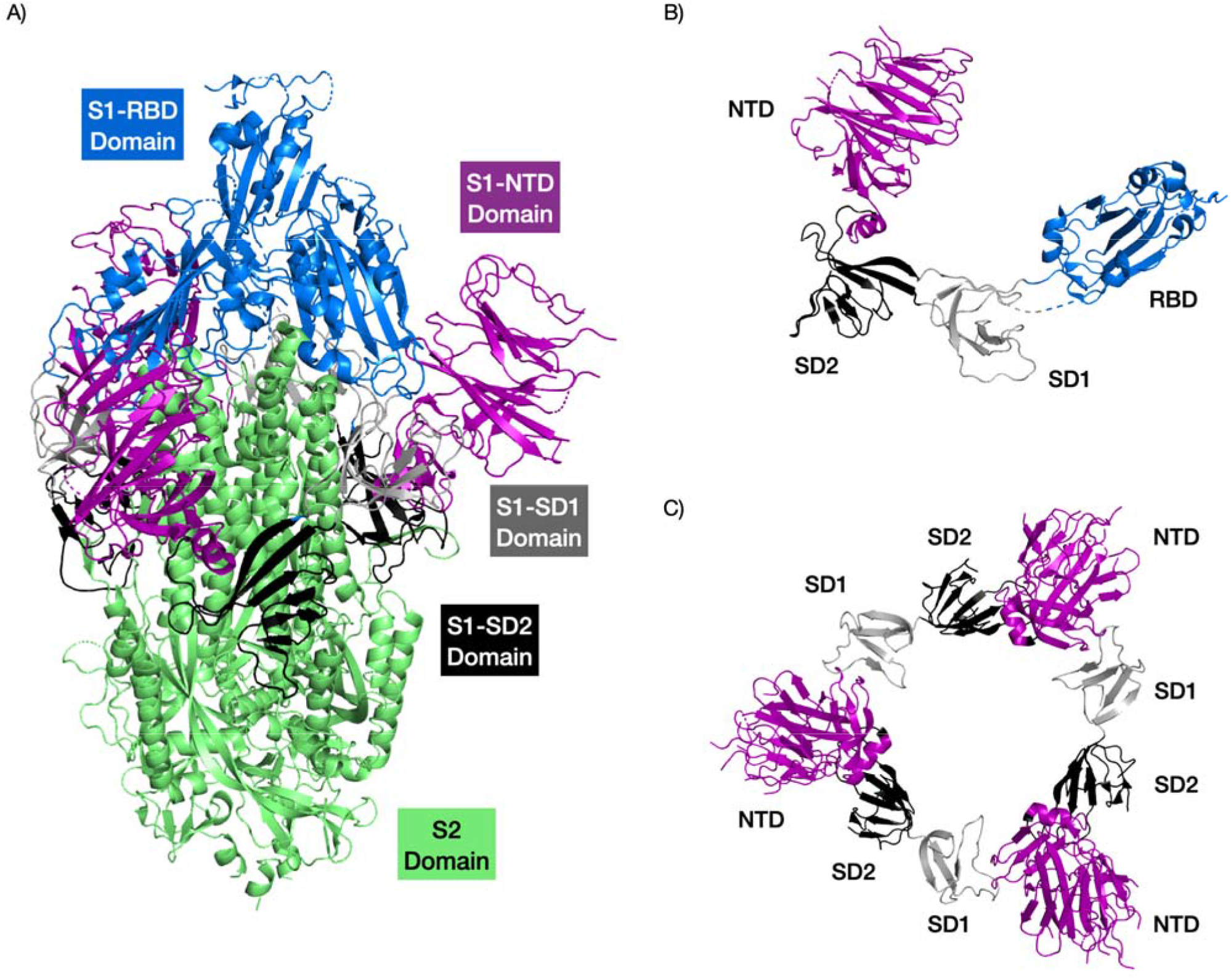
Structure of the SARS-CoV2 Spike protein highlighting the different domains. The RBD domain is coloured in blue, the NTD domain in purple, the SD1 domain in grey and the SD1 domain in black. The trimer complex is shown in (A). The S1 region of a Spike protein monomer is shown in (B). The interactions of the NTDs with SD1 and SD2 domains of the Spike protein trimer are shown in (C). PDB structure 6VSB.

Structure analyses of the 4 BCoVs that are in the same lineage (B) (see above) as SARS-CoV-2, showed that the NTD domains are all assigned to the same evolutionary superfamily (2.60.120.960) in the in-house CATH domain structure classification (Sillitoe et al., 2021). This superfamily falls in the fold group (2.60.120) containing the human sugar-binding protein Galectin-3. To determine how conserved this domain is across these 4 BCoVs and SARS-CoV-2 are, and identify any variable regions, we performed a structure superposition using our in-house protein structure alignment tool SSAP (see methods). Overall, the structures are very similar (see Figure 3) with an average SSAP of 86 (range is 0 to 100) suggesting a high global conservation of this domain across these BCoVs. However, whilst a large core of the domain is highly conserved, there are extensions in some of the loops. The variable nature of these loop regions across multiple BCoV NTD domains was also identified in a multiple sequence alignment (Garry et al., 2021), but the authors did not perform structural analyses to assess their proximity to putative sugar-binding regions. However, there is some evidence in other BCoVs that these changes may improve affinity for sugars on the host cell surface and therefore increase the affinity of the virus for the specific host cell (Hulswit et al., 2019; Qing et al., 2020).

**Figure 3.**
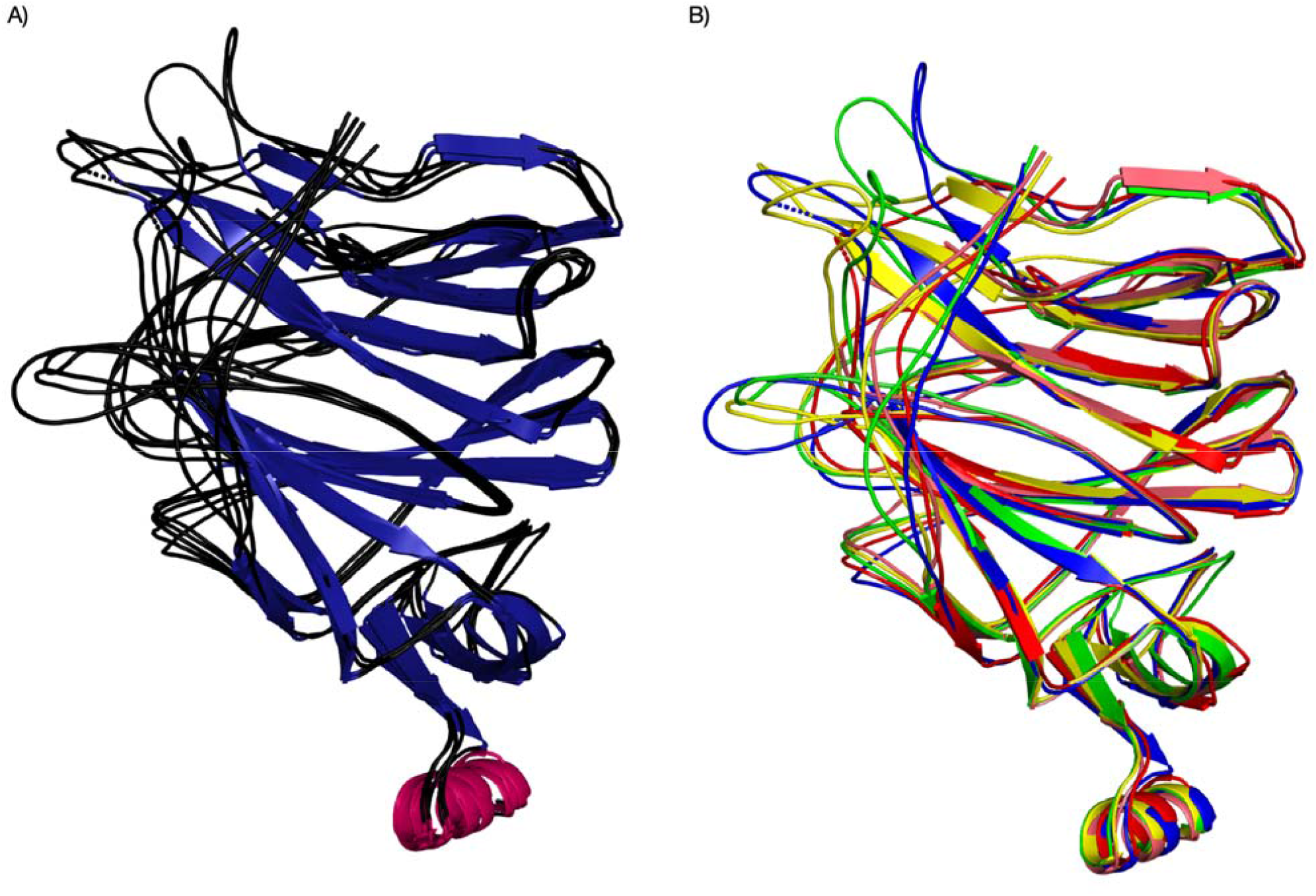
Structure superposition of Cov NTDs (SARS-CoV, PDB ID 6ACC, Pangolin CoV GX, PDB ID 7CN8, Pangolin-CoV-GD, PDB-ID 7BBH, Bat-Cov-RaTG13, PDB ID: 7CN4, SARS-CoV-2, PDB ID 7C2L). The structures are coloured based on their secondary structure components (A) and proteins (different species/variants) (B).

### 3.3 Analysis of pockets in the NTD domain

As mentioned above, there is evidence of at least three pockets in SARS-CoV-2 (Awasthi et al., 2020; Fantini et al., 2020; Baker et al., 2021; Di Gaetano et al., 2021), two of which are known to bind sialic acid/sugars. To confirm and further characterise these and search for additional pockets we used cavity detection and docking tools. Analysing the structure of the SARS-CoV-2 NTD domain using CavityPlus (Xu et al., 2018), we identified the known sugar binding pocket shared with Galectin-3 (pocket 1, Figure 4). Two other cavities were confirmed (pocket 2 (red) and 3 (blue in Figure 4) that share, through a common loop, a reported conserved sugar-binding motif (comprising 7 residues - G72, T73, N74, G75, T76, K77 and R78). Furthermore, in agreement with McCallum et al. (2021) we observe that the antigenic NTD supersite coincides with sugar-binding pockets 1 and 2. Finally, a fourth, previously undefined pocket was detected (pocket 4, colour cyan) shown in figure 4 below.

**Figure 4.**
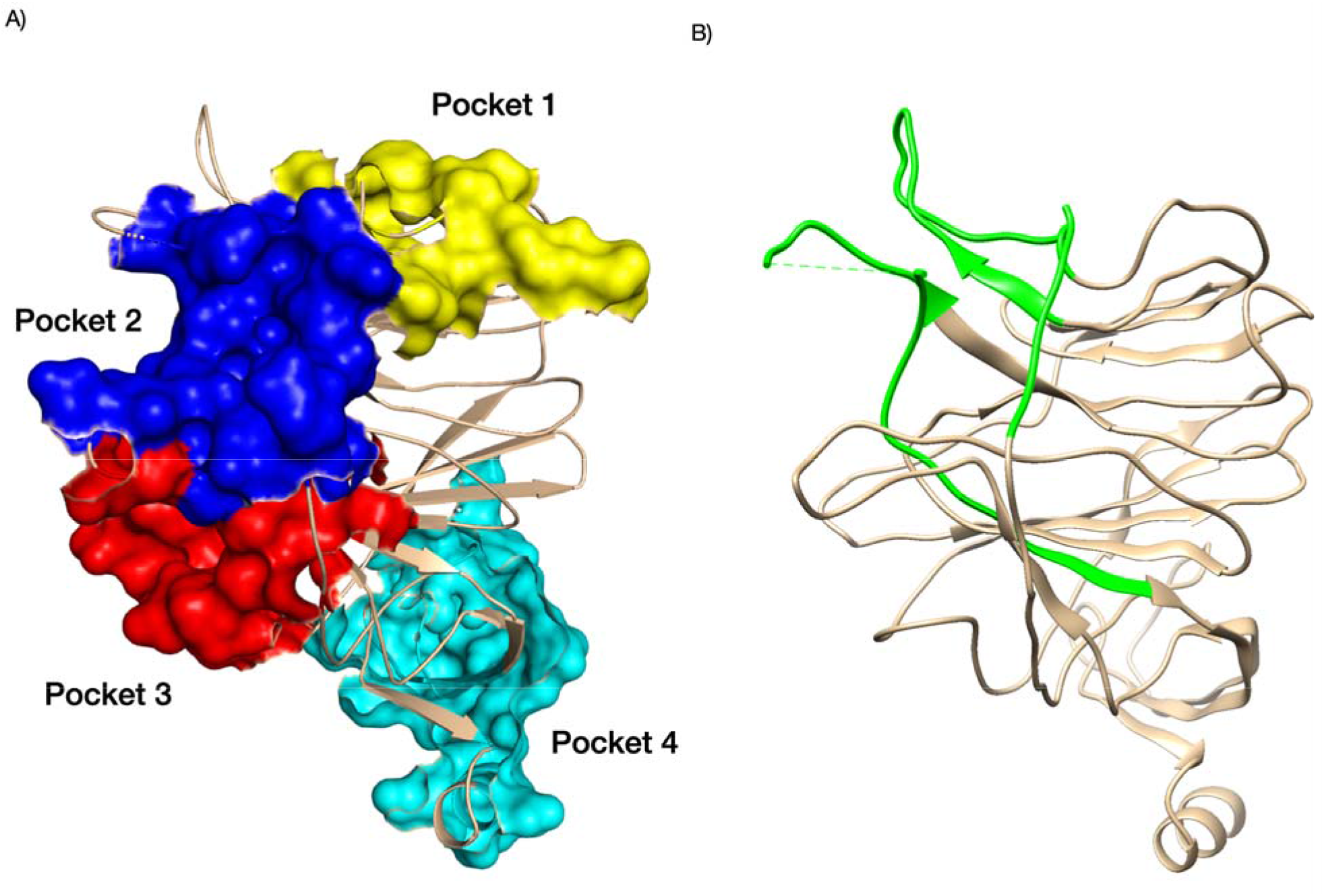
Pockets identified in NTD domain of SARS-CoV-2 (A) and antigenic NTD supersite of SARS-CoV-2 shown in green (B). For (A), we coloured pocket 1 in yellow, pocket 2 in blue, pocket 3 in red and pocket 4 in cyan. PDB structures 7C2L. Supporting references: Behloul et al., 2020; Fantini et al., 2020; Baker et al., 2021; Di Gaetano et al., 2021; McCallum et al., 2021.

Supplementary Table 4 shows the SARS-CoV-2 NTD residues that form the pockets detected by our study. Some of these (pocket1 and 2) are thought to interact with sugars (e.g. sialic acid) (Fantini et al., 2020; Behloul et al., 2020, Baker et al., 2021). Docking studies performed by us (see Section 3.6 below) also suggest pocket 3 and 4 may bind sialic acid. Studies that previously identified or commented on these pockets are cited in the table.

### 3.4 Multiple sequence alignment of the BCoVs reveals evolutionary hotspots

Garry et al. performed a detailed sequence analysis of multiple BCoVs and strains of SARS-CoV-2 and demonstrated the suitability of this approach to highlight regions varying across the BCoVs. We have expanded this alignment to 31 BCoVs and used a structure guided approach to ensure accurate alignment of more distant BCoVs (Figure 5). Garry et al (2021) observed that lineage indels are a common feature of the Spike protein in BCoVs, with seven major loops most frequently affected. Five out of seven loops involve the NTD. We aligned the sequences of the 31 BCoVs included in the phylogenetic analysis using CLUSTAL-OMEGA, excluding the MERS-CoV and HCoV-OC43 BCoVs as they are from a different lineage and disrupt the alignment. Since we had structural data for SARS-CoV-2, we exploited this data to produce a structure-based multiple sequence alignment (MSA), using ESPript.

**Figure 5.**
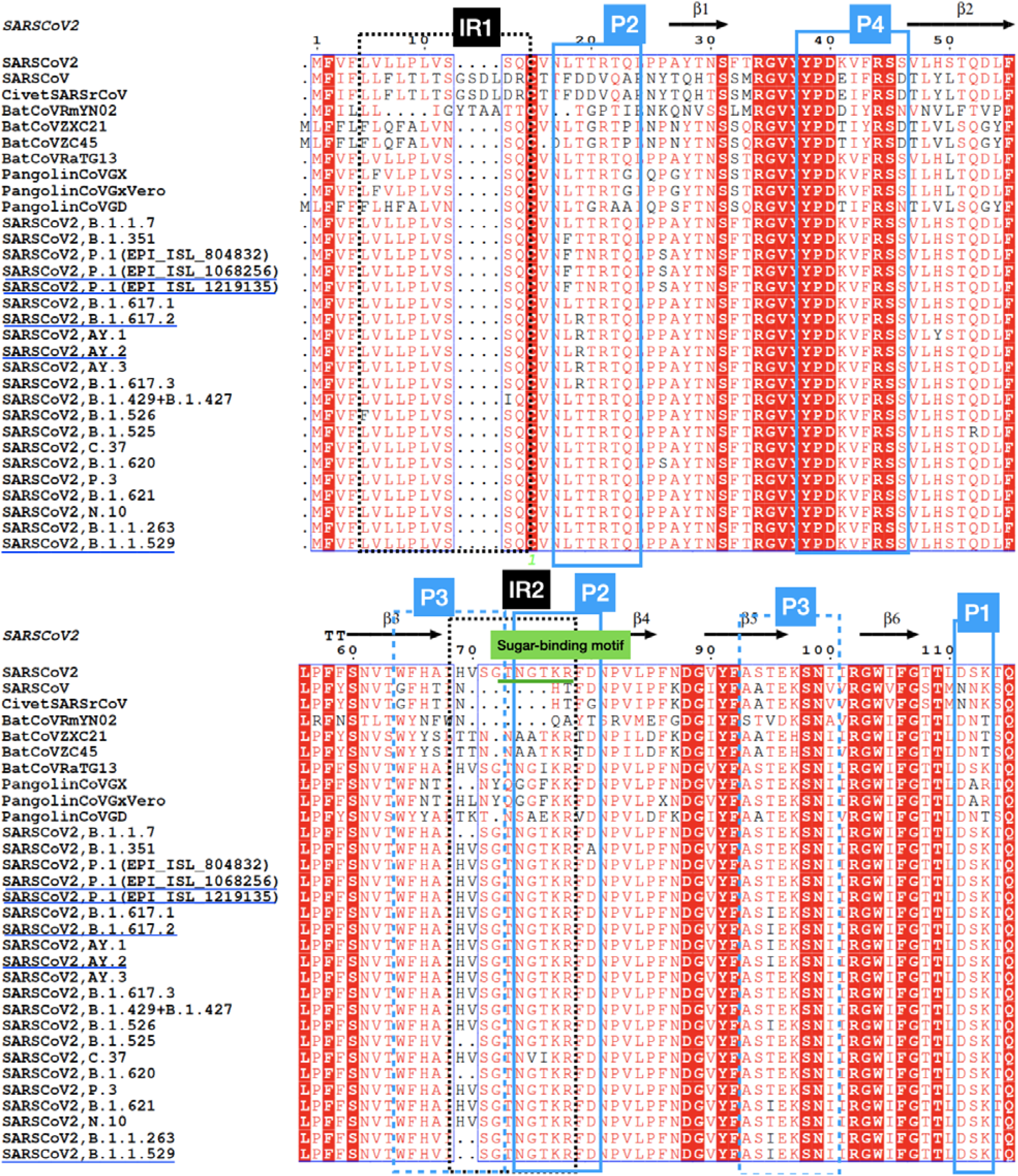
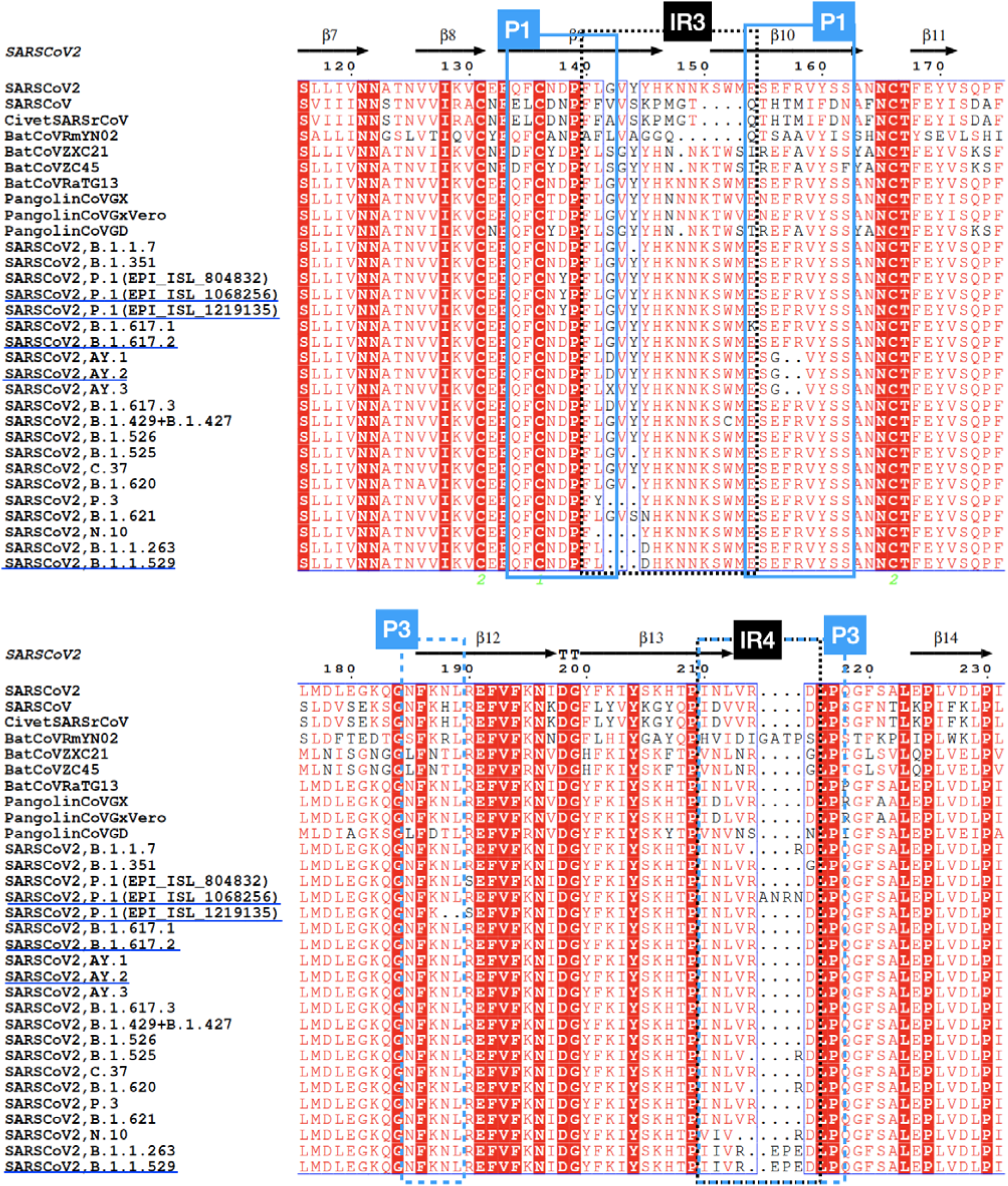
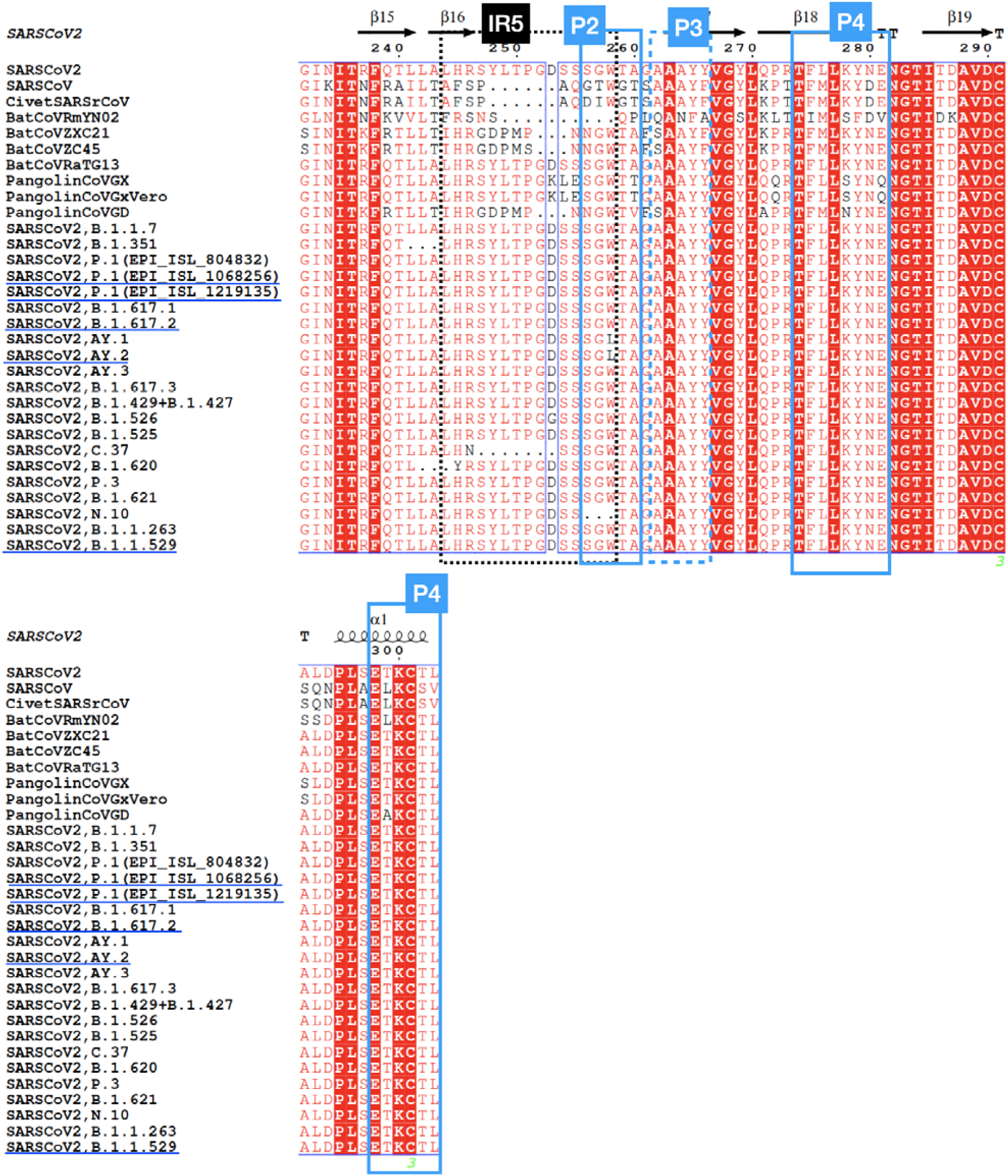
Multiple sequence alignment of BCoV NTDs. The Indel region (IRs), sugar-binding motif and known/putative sugar-binding pockets (P) are highlighted. Indel regions (IR1-IR5), pockets (P1-P4). VOCs having variants that affect binding energy are underlined (i.e. P.1 Gamma variant, B.1.617.2 Delta variant, AY.2 Delta Plus variant, B.1.1.529 Omicron variant)

Whilst residues in most regions of the protein are highly conserved (ScoreCons values > 80), the residues contributing to sugar binding pocket 1 are somewhat less conserved (pocket 1 – average ScoreCons of 70) and even more so for pocket 2 (average ScoreCons of 61). Pocket 2 has changed in some BCoVs (SARS-CoV, civet SARS-CoV and BatCoV-RmYN02). Cheng et al. also compared BCoVs NTD structures and found pocket 2 of SARS-CoV to be less well-defined with the loops not giving such a well-formed pocket. Our docking studies (see Section 3.6 below) suggest that pocket 3 is also able to bind sialic acid/sugars. This pocket is also quite variable with average ScoreCons values of 73. However, pocket 4 is very conserved relative to the other pockets with an average ScoreCons value of 87.

The proximity of the indel regions to residues in these known and putative sugar-binding pockets is shown in Table 1, below. Most are close to pockets 2 and 3. Indel region 2 (close to pockets 2 and 3), indel region 3 and indel region 5 (close to pocket 3) are the most variable regions across these BCoVs.

**Table 1.** Proximity of indel regions to residues in the sugar binding pockets. If the indel region lies close to multiple pockets (<5Å), minimal residue distances to all pockets are provided.

| Indel region | Minimal residue distance to known and putative sugar binding pockets (Å) |
| --- | --- |
| Indel region 1 | 1.86 (Pocket 1)<br>1.33 (Pocket 2) |
| Indel region 2 | 1.33 (Pocket 2)<br>1.33 (Pocket 3) |
| Indel region 3 | 7.32 (Pocket 1) |
| Indel region 4 | 1.31 (Pocket 3) |
| Indel region 5 | 1.31 (Pocket 2) |

We see that much of the indel innovation between lineages occurs in loop regions associated with the NTD binding pockets 2 and 3 suggesting that these binding pockets are hotspots for viral evolution (Figure 6). Indeed, most of the indels occur on this side of the NTD domain (See Figure 6) and seem to target the sugar-binding pockets, possibly tuning the interactions with sugars such as sialic acids.

**Figure 6.**
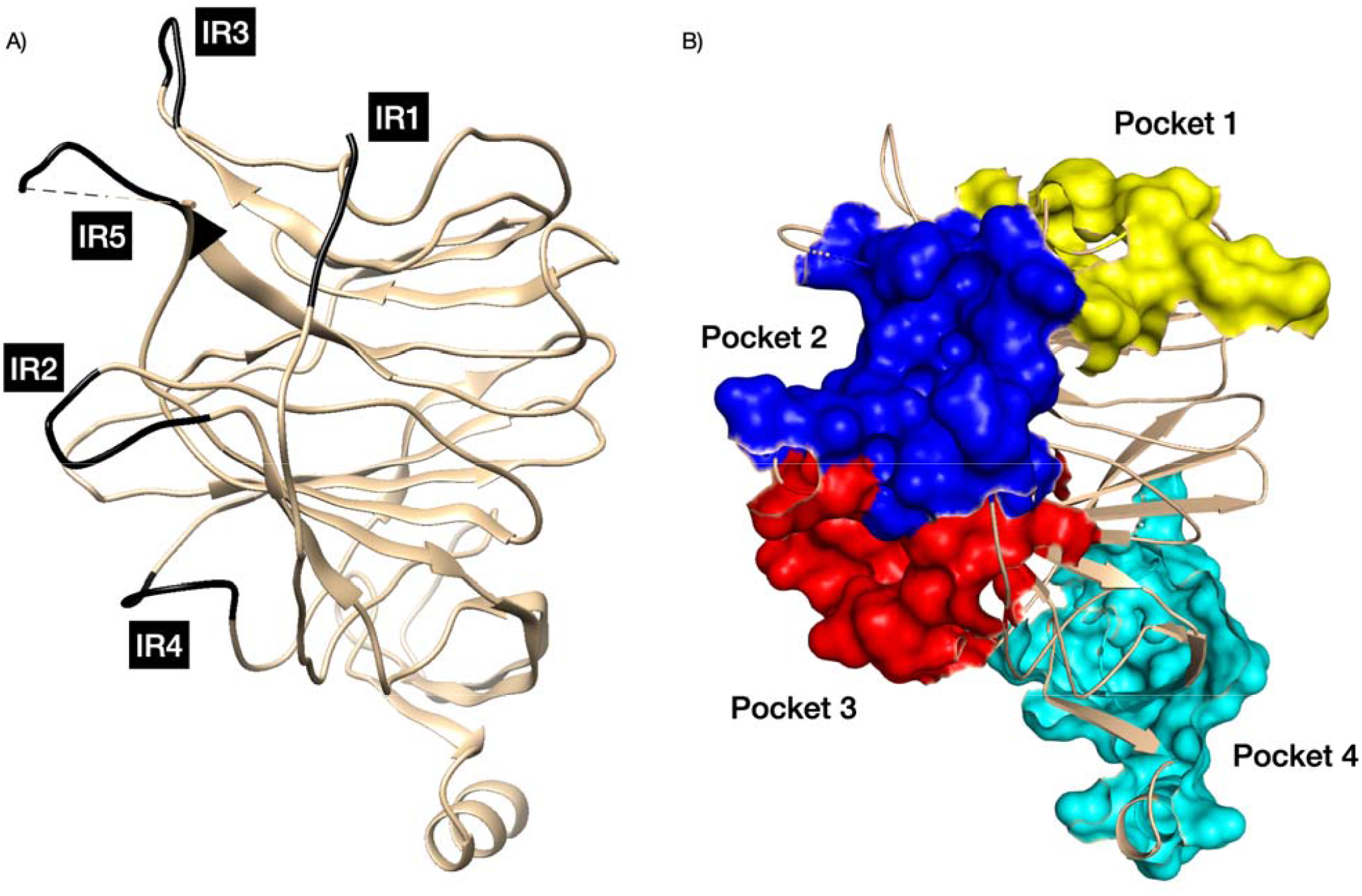
Indel regions and binding pocket of SARS-CoV-2 NTD. (A) Indel regions (IR) were identified using the MSA. (B) Highlighting the sugar-binding pockets, we coloured pocket 1 in yellow, pocket 2 in blue, pocket 3 in red and pocket 4 in cyan. PDB structures 7C2L.

### 3.5 Structural analysis of evolution of the sugar-binding pockets

We analysed the structural features observed in different BCoV domains in the CATH superfamily containing NTD domains (see Figure 7). We also compared against the human Galectin-3 protein which is in the same fold group as this superfamily. It can be seen that loop insertions in the BCoVs near the sugar-binding pocket 1 appear to enhance the contact of the protein with the ligand and the contacts increase going from human Galectin-3 to HCoV-OC43 (a BCoV that emerged in the 19th century) up to SARS-CoV-2.

**Figure 7.**
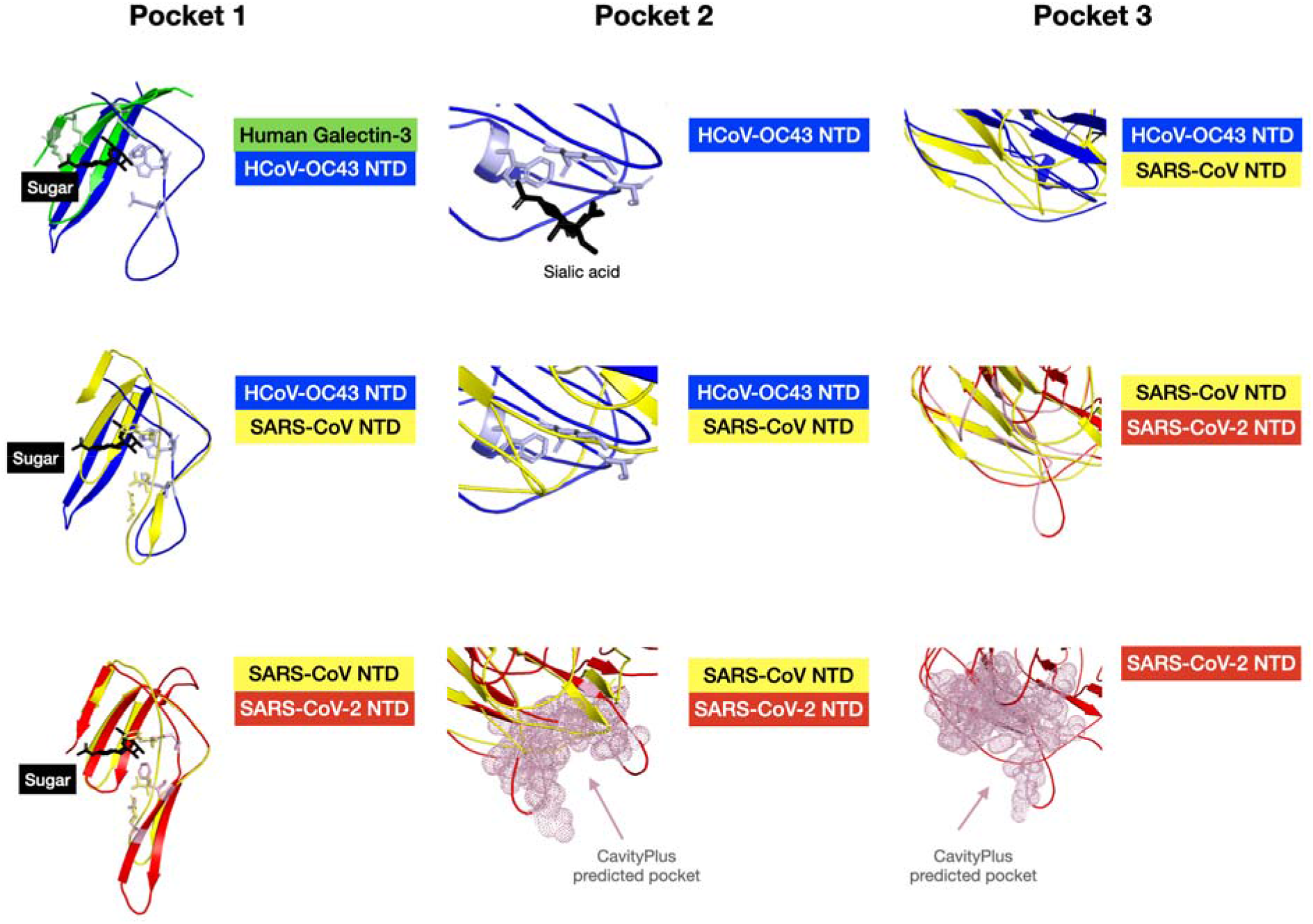
Evolution of the sugar-binding pocket from human Galectin-3 to SARS-CoV-2 and emergence of other sugar-binding regions. We only show the sugar-binding regions. We coloured the human Galectin-3 NTD structure in green, hCoV-OC32 NTD structure in blue, SARS-CoV NTD structure in yellow and SARS-CoV-2 NTD structure in red. PDB structures 1A3K,7C2L,6ACC, 6NZK respectively.

**Figure 8.**
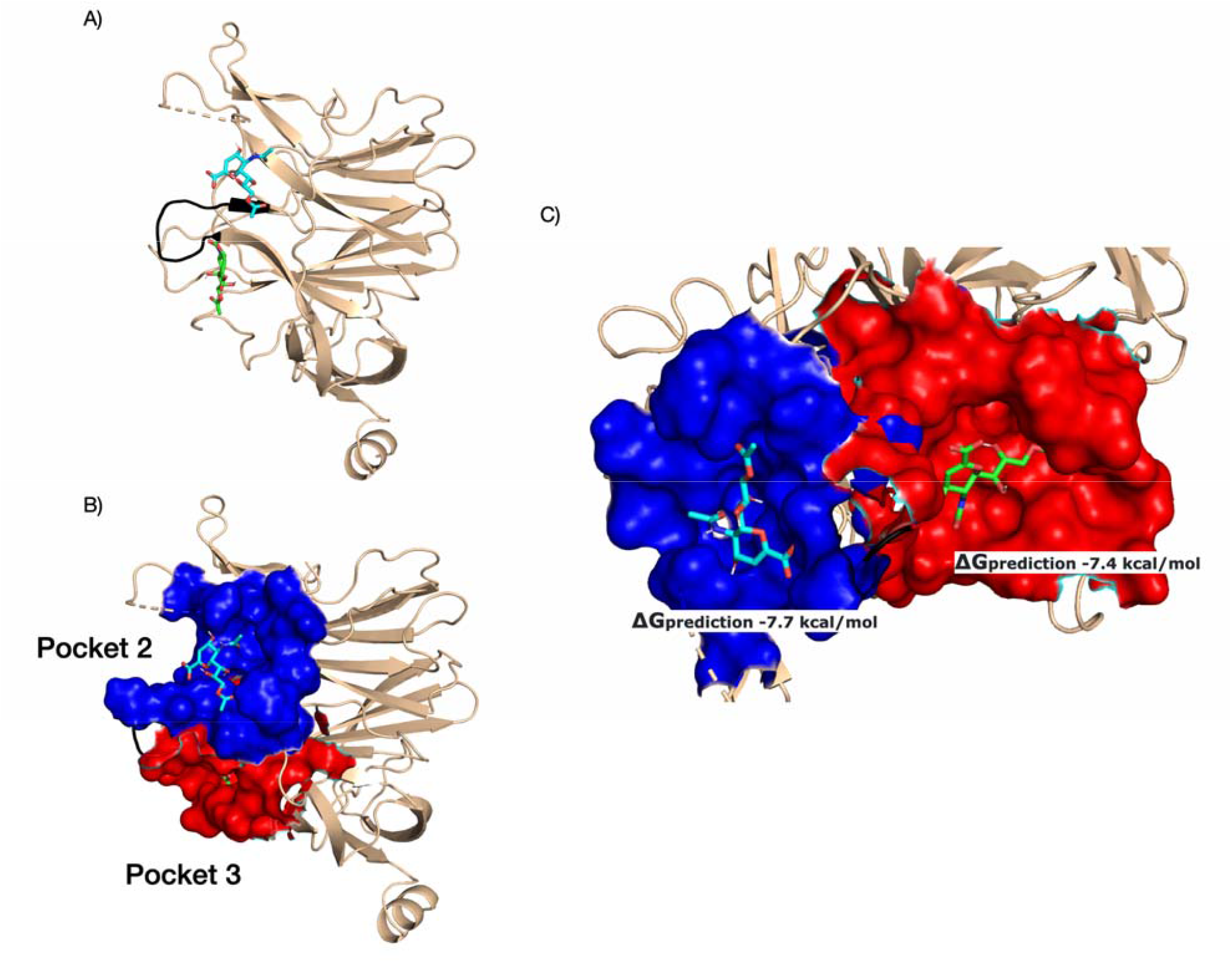
Sialic acid docked into two putative cavities in the SARS-CoV2 NTD domain. A) the sugar-binding motif is shown in black, following the same orientation as in the other figures. B) cavities were identified by CavityPlus and are shown using surface representation and coloured blue (pocket 2) and red (pocket 3) respectively. C) the structure is oriented to show both cavities and we give the PRODIGY predicted binding energy.

As mentioned in the introduction, Behloul and co-workers found a GTNGTKR motif (located within a different pocket in the BCoVs (Pocket 2) suggesting that this region may also bind sugars (Behloul et al., 2020). Human Galectin-3 does not bind sugar at this pocket (Johannes et al., 2018) and we also see loop extensions when comparing SARS-CoV-2 to human Galectin-3 in the region around pocket 2 (see Figure 7).

Loops in the region of pocket 2 seem to be highly structurally variable and this pocket is less well defined in SARS-CoV. Comparing the structures and sequences of HCoV-OC43 and SARS-CoV-2 we see strong innovation in pocket 2 in SARS-CoV-2 suggesting alterations in this pocket were important for the evolution of this virus. Similarly, structural analyses comparing pocket 3 across the coronaviruses suggests that this pocket can be highly variable too, and it is also found to be well defined in SARS-CoV-2 offering another potential sialic acid binding site. We used docking studies to explore and contrast the ability of the different pockets in SARS-CoV-2 to bind sialic acid.

### 3.6 Docking analyses of sialic acid in pockets of SARS-CoV-2

Previous studies had indicated that pocket 1, which is similar to the sugar binding pocket in Galactin-3, could bind sugar (Fantini et al., 2020) and that pocket 2 could possibly bind sugars. For example, a structure of a BCoV (HCoV-HKU23 – which has 25% sequence identity to SARS-CoV-2) with sialic acid bound in pocket 2 has been solved (Tortorici et al., 2019). Therefore, whilst there is some indication, including from MD studies (Awasthi et al., 2020; Fantini et al., 2020; Behloul et al., 2020; Baker et al., 2021) that SARS-CoV-2 can bind sialic acid in pockets 1 and 2, further computational analyses were used to explore and contrast the binding of sialic acid to all pockets in the NTD domain.

We first determined the binding energy of sialic acid bound to pocket 2 of HCoV-HKU23 using the PRODIGY score of HADDOCK and obtained a value of -7.1 kcal/mol. The LigPlot for the structure-ligand complex is shown in Supplementary Figure 3. We then used HADDOCK to dock sialic acid into all four of the pockets in the experimental structure of the NTD domain of SARS-CoV-2, and calculated the PRODIGY predicted binding energy of sialic acid. The results suggest that sialic acid binds most strongly to pockets 2 and 3 (see Table 2). Awasthi and co-workers also demonstrated the ability of other sialic acids (such as 5-N-acetyl neuraminic acid (Neu5Ac), α2,3-sialyl-N-acetyl-lactosamine (2,3-SLN), α2,6-sialyl-N-acetyl-lactosamine (2,6-SLN), 5-N-glycolyl neuraminic acid (Neu5Gc), and sialyl LewisX (sLeX)) to bind into pocket 2 of SARS-CoV-2 using in silico molecular docking (2020).

**Table 2.** PRODIGY predicted binding energy of sialic acid to NTD pockets of SARS-CoV and SARS-CoV-2

| Pocket | PRODIGY<br>predicted binding<br>energy (Pocket<br>1) | PRODIGY<br>predicted binding<br>energy (Pocket<br>2) | PRODIGY<br>predicted binding<br>energy (Pocket<br>3) | PRODIGY<br>predicted binding<br>energy (Pocket<br>4) |
| --- | --- | --- | --- | --- |
| SARS-CoV | -5.5 kcal/mol | -6.4 kcal/mol | -6.6 kcal/mol | -6.4 kcal/mol |
| SARS-CoV-2 | -6.6 kcal/mol | -7.7 kcal/mol | -7.4 kcal/mol | -6.6 kcal/mol |

Since pockets 2 and 3 show the most variability across BCoVs and bind sialic acid more strongly than the other pockets we subsequently used LigPlot+ to further analyse the interactions of sialic acid with residues in these pockets. For pocket 2 (see Supplementary Figure 2), 3 residues of the sugar binding motif (T76, K77 and R78) are involved in binding with sialic acid. K77 forms a hydrogen bond and hydrophobic interaction with sialic acid, while T76 and R78 form hydrophobic interactions with the compound. For pocket 3, LigPlot showed no polar interactions of the ligand with residues in the sugar-binding motif. Yet, we also observe hydrogen bonds and Van der Waals interactions between sialic acid and the NTD.

#### 3.6.1 Comparisons with other BCoVs

Since pocket 2 and 3 are the most variable and less frequently studied pockets, we compared these putative sialic binding pockets in SARS-CoV-2 to SARS-CoV and Pangolin-CoV-GX. We used HADDOCK to dock sialic acid into the experimental structures of the NTD domains of these BCoVs. We selected the same binding residues as pocket 2 and 3 in SARS-CoV-2. None of these proteins binds sialic acid as strongly as SARS-CoV-2 (see Table 3). This result is not surprising as previous structural analysis by Cheng et al. found pocket 2 of SARS-CoV to be less-defined than other BCoVs (2019). We also compared the LigPlots for sialic acid docked to SARS-CoV and Pangolin-CoV-GX with those for SARS-CoV-2 (See Supplementary Figure 3 and Supplementary Figure 4). For both pockets, we observe more interactions with sialic acid and more hydrogen bonds formed in SARS-CoV-2 NTD than SARS-CoV NTD. The number of sialic acid interacting residues for Pangolin-CoV-GX and SARS-CoV-2 are similar.

**Table 3.** PRODIGY predicted binding energy of sialic acid to NTD pocket 2 and pocket 3 of selected BCoVs

| Coronavirus | PRODIGY predicted binding energy (Pocket 2) | PRODIGY predicted binding energy (Pocket 3) |
| --- | --- | --- |
| SARS-CoV | -6.4 kcal/mol | -6.6 kcal/mol |
| Pangolin-CoV-GX | -7.3 kcal/mol | -7.2 kcal/mol |
| SARS-CoV-2 | -7.7 kcal/mol | -7.4 kcal/mol |

### 3.7 Analysis of recent variants of concern/interest in SARS-CoV-2

In order to assess the possible impact of recent variants of concern/interest in relation to these sugar binding pockets, we performed molecular docking of sialic acid into all 3 pockets using the original Wuhan-1 strain and assessed the impacts on the binding for the various mutations in the VOCs and VOIs. We did not consider pocket 4 as this is highly conserved across BCoVs and no VOC variants occurred in this pocket. We list all the NTD mutations/insertions found in the different VOCs/VOIs in Supplementary Table 5. Many of the mutations/insertions lie close to the 3 known and putative sugar binding pockets discussed in this and related studies.

We report the PRODIGY binding energy for the individual NTD mutations in Supplementary Table 6. Overall, most of the mutations in the VOCs and VOIs were not predicted to drastically alter or enhance sialic acid binding. However, two mutations, one observed in pocket 1: E154K (found in Kappa variant) and the other observed in pocket 3: T95I (found in Kappa, Delta, Delta Plus (AY.1, AY.2) Iota, Mu and B.1.1.263 variants) were predicted to increase binding affinity (≥0.5 kcal/mol changes), as compared to the original Wuhan-1 strain of SARS-CoV-2. See Supplementary Figure 6 and Supplementary Figure 7 for LigPlots.

We also examined the impact on binding energies for NTD domains in different strains, when all the mutations and indels in the domain were considered (see Table 4). For pocket 1, the binding of the B.1.617.2 Delta strain, AY.2 Delta plus strain and two of the Gamma P.1 strains to sialic acid are stronger than the original Wuhan-1 strain (an increase of 0.5 kcal/mol) (See Supplementary Figure 8 for LigPlot). For pocket 2, there is no significant difference among the SARS-CoV-2 strains. However, for pocket 3 we see a significant increase in binding energy, compared to the original SARS-CoV-2, in the following strains of SARS-CoV2: B.1.617.3, B.1.429+B.1.427 (Epsilon variant), B.1.526 (Iota variant), C.37 (Lambda variant), P.3 (Theta variant), N.10, B.1.1.263 and B.1.1.529 (Omicron variant) (See Supplementary Figure 9 for LigPlots). However, it is important to note that these predictions need to be verified by careful experimental work to confirm these effects.

**Table 4.**
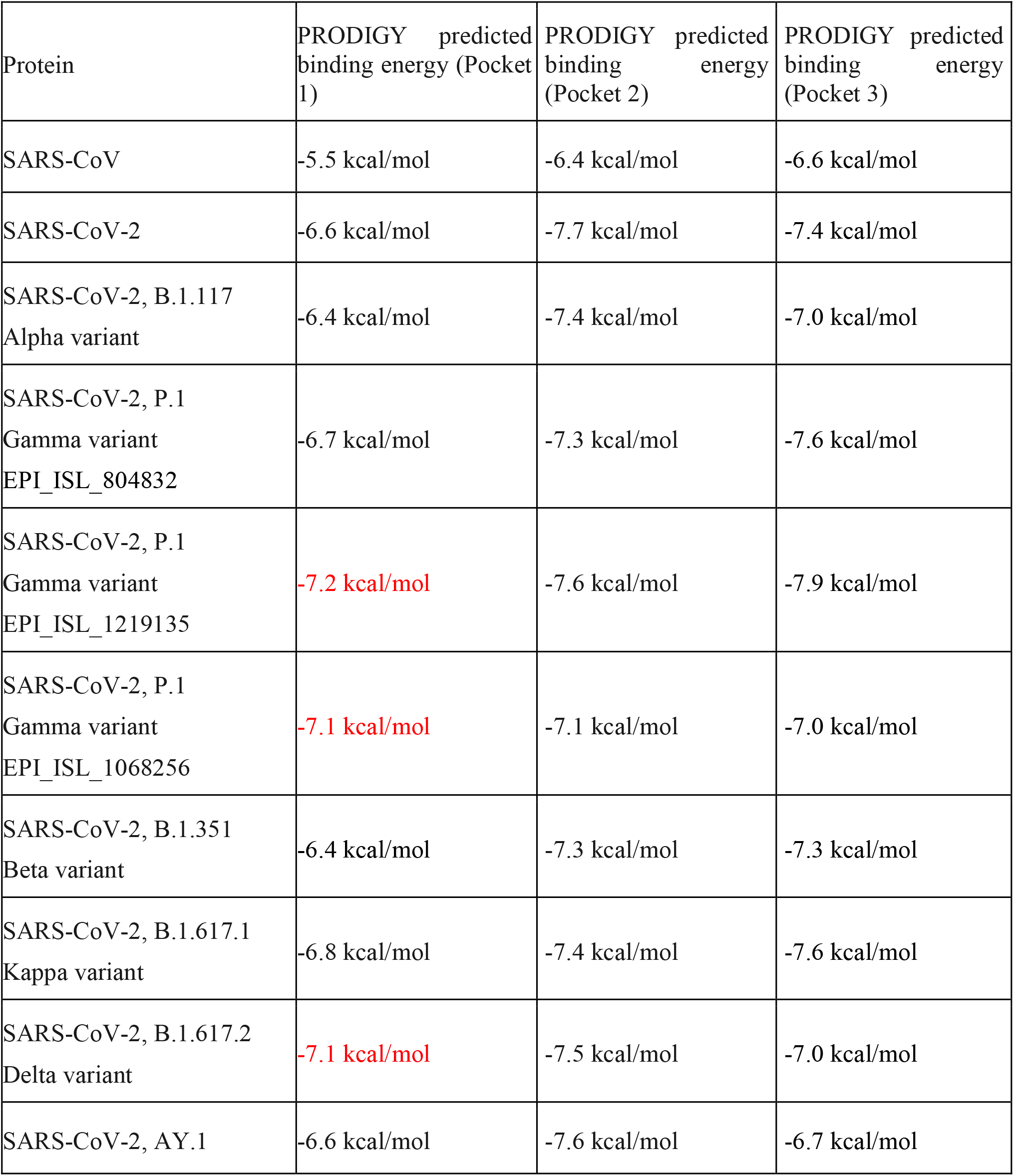
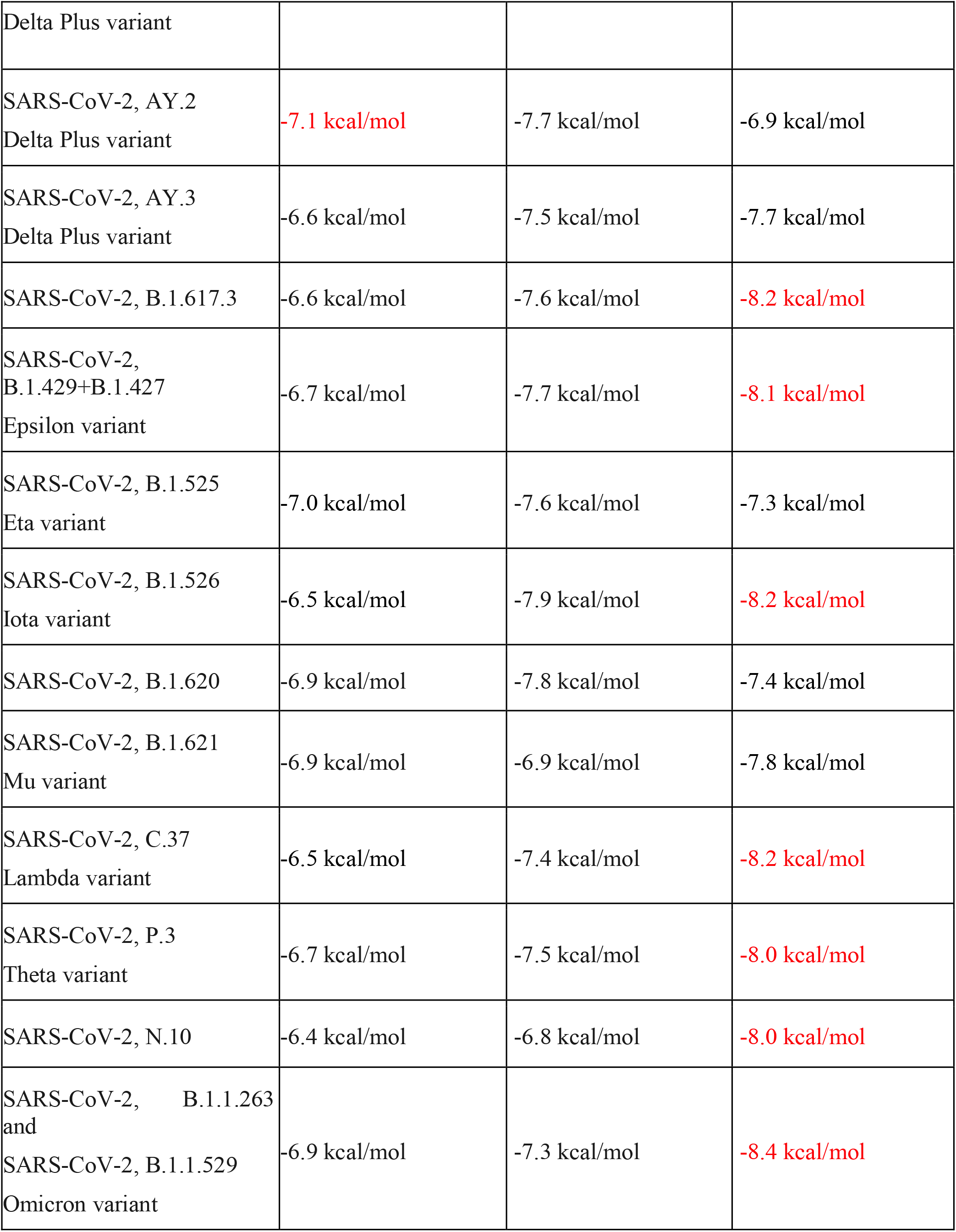
PRODIGY predicted binding energy of sialic acid to SARS-CoV NTD pockets. Strain with an increase binding affinity of ≥0.5 kcal/mol as compared to the original Wuhan-1 strain of SARS-CoV-2 is coloured red.

| Protein | PRODIGY predicted binding energy (Pocket 1) | PRODIGY predicted binding energy (Pocket 2) | PRODIGY predicted binding energy (Pocket 3) |
| --- | --- | --- | --- |
| SARS-CoV | -5.5 kcal/mol | -6.4 kcal/mol | -6.6 kcal/mol |
| SARS-CoV-2 | -6.6 kcal/mol | -7.7 kcal/mol | -7.4 kcal/mol |
| SARS-CoV-2, B.1.117<br>Alpha variant | -6.4 kcal/mol | -7.4 kcal/mol | -7.0 kcal/mol |
| SARS-CoV-2, P.1<br>Gamma variant<br>EPI_ISL_804832 | -6.7 kcal/mol | -7.3 kcal/mol | -7.6 kcal/mol |
| SARS-CoV-2, P.1<br>Gamma variant<br>EPI_ISL_1219135 | -7.2 kcal/mol | -7.6 kcal/mol | -7.9 kcal/mol |
| SARS-CoV-2, P.1<br>Gamma variant<br>EPI_ISL_1068256 | -7.1 kcal/mol | -7.1 kcal/mol | -7.0 kcal/mol |
| SARS-CoV-2, B.1.351<br>Beta variant | -6.4 kcal/mol | -7.3 kcal/mol | -7.3 kcal/mol |
| SARS-CoV-2, B.1.617.1<br>Kappa variant | -6.8 kcal/mol | -7.4 kcal/mol | -7.6 kcal/mol |
| SARS-CoV-2, B.1.617.2<br>Delta variant | -7.1 kcal/mol | -7.5 kcal/mol | -7.0 kcal/mol |
| SARS-CoV-2, AY.1 | -6.6 kcal/mol | -7.6 kcal/mol | -6.7 kcal/mol |
| Delta Plus variant |  |  |  |
| SARS-CoV-2, AY.2<br>Delta Plus variant | -7.1 kcal/mol | -7.7 kcal/mol | -6.9 kcal/mol |
| SARS-CoV-2, AY.3<br>Delta Plus variant | -6.6 kcal/mol | -7.5 kcal/mol | -7.7 kcal/mol |
| SARS-CoV-2, B.1.617.3 | -6.6 kcal/mol | -7.6 kcal/mol | -8.2 kcal/mol |
| SARS-CoV-2,<br>B.1.429+B.1.427<br>Epsilon variant | -6.7 kcal/mol | -7.7 kcal/mol | -8.1 kcal/mol |
| SARS-CoV-2, B.1.525<br>Eta variant | -7.0 kcal/mol | -7.6 kcal/mol | -7.3 kcal/mol |
| SARS-CoV-2, B.1.526<br>Iota variant | -6.5 kcal/mol | -7.9 kcal/mol | -8.2 kcal/mol |
| SARS-CoV-2, B.1.620 | -6.9 kcal/mol | -7.8 kcal/mol | -7.4 kcal/mol |
| SARS-CoV-2, B.1.621<br>Mu variant | -6.9 kcal/mol | -6.9 kcal/mol | -7.8 kcal/mol |
| SARS-CoV-2, C.37<br>Lambda variant | -6.5 kcal/mol | -7.4 kcal/mol | -8.2 kcal/mol |
| SARS-CoV-2, P.3<br>Theta variant | -6.7 kcal/mol | -7.5 kcal/mol | -8.0 kcal/mol |
| SARS-CoV-2, N.10 | -6.4 kcal/mol | -6.8 kcal/mol | -8.0 kcal/mol |
| SARS-CoV-2, B.1.1.263<br>and<br>SARS-CoV-2, B.1.1.529<br>Omicron variant | -6.9 kcal/mol | -7.3 kcal/mol | -8.4 kcal/mol |

### 3.8 Pocket 1, 2 and 3 are variable across BCoVs but structural analyses reveal the fourth pocket is highly conserved and potentially druggable

Structural superpositions of the known (PDB) structures of BCoV NTD domains are shown in Figure 9B. We also modelled the structures of all BCoVs classified in the same superfamily in CATH, using AlphaFold2. Only good quality models (predicted IDDT score >75, See Supplementary Table 3 for details) were used and we superposed these to assess the conservation of the structures (see Figure 9B). Overall, the conservation of the structures is high with an average SSAP of 87 (out of a maximum score of 100). We also calculated the structure conservation for the pockets. Pocket 1 is conserved with an average SSAP of 83.50. There have been attempts to design drugs for this pocket and chloroquine and hydroxychloroquine have been tested and shown to bind in this pocket [Fantini et al (2020)]. These drugs have been used for the treatment of malaria but are not being used for Covid-19. On the other hand, the structural conservation for pockets 2 and 3 is low, with average SSAP scores of 65-73.

**Figure 9.**
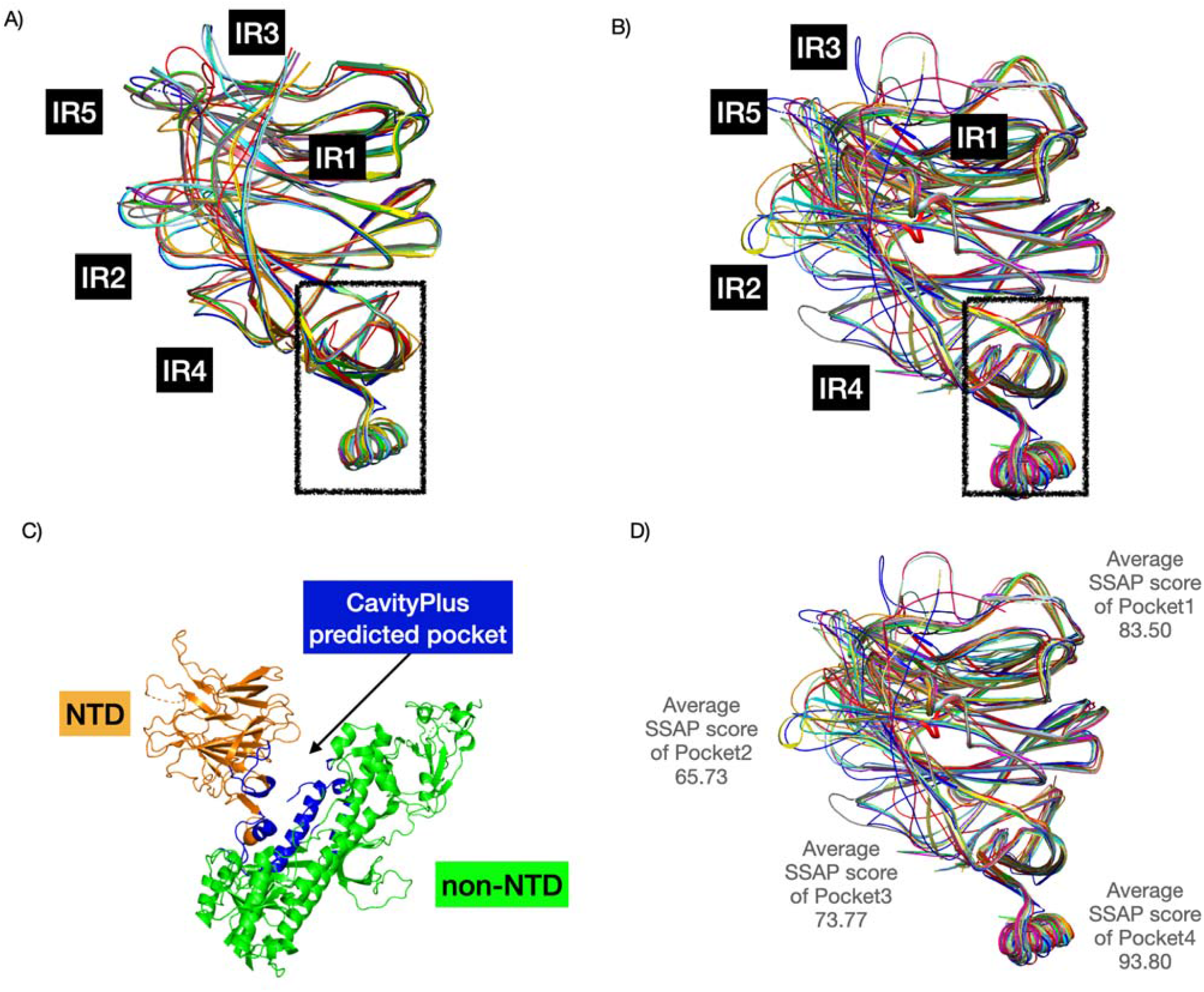
Structure superposition of CoV NTDs. A) comprises the 16 BCoV NTDs in the multiple sequence alignment (see Figure 5). B) constitutes 24 BCoV NTDs from the CATH family. For A) and B), we used known structures and structural models (built using AlphaFold2). We found a highly conserved pocket (in the box) (score of 1838 - highly positive DrugScore constitutes high druggability) that could be a good pan coronavirus drug target. C) demonstrates this pocket predicted by CavityPlus in blue. D) We computed the structural conservation of pockets by calculating the average SSAP score.

However, analysis by CavityPlus identified a fourth pocket druggable pocket (CavityPlus Drug Score of 1838, a highly positive DrugScore suggests high druggability), which binds sialic acids less strongly than the other pockets. The structural conservation of this pocket is very high (average SSAP score of 93.80). The high structural conservation across BCoVs and the druggability of this pocket suggest that it could be a good pan coronavirus drug target.

## 4 Discussion

Several experimental studies have reported that BCoVs use sialic acid-containing receptors for entry into the host cell (Tortorici et al., 2019; Peng et al., 2021). In the case of SARS-CoV-2, sequence and structure based studies reported critical residues involved in putative sugar binding pockets (pockt 1 and 2) (Behloul et al., 2020; Fantini et al., 2020; Baker et al., 2021).

We have built on these studies and examined all known and putative binding pockets in the NTD domain of SARS-CoV-2 and related coronaviruses. Whilst pockets 1 and 2 previously had detailed structural analyses including MD and docking (Awasthi et al., 2020; Fantini et al., 2020), suggesting sialic acid binding, pocket 3 whilst detected (Di Gaetano et al., 2021) had not been characterised for sialic acid binding. Furthermore, to our knowledge pocket 4 had not been reported by other groups. We found pocket 2 and 3 to bind sialic acid more strongly than pocket 1. Pocket 3 binds sialic acid as strongly as pocket 2 and like pocket 2 is highly variable across BCoVs. Unlike other studies on these pockets, our study also includes comprehensive analyses of Variants of Concern (VOCs) and Variants of Interest (VOIs) in these pockets. When considering all changes in the VOC, some significant changes in binding energy were linked to pocket 3 (B.1.617.3, B.1.429+B.1.427 (Epsilon variant), B.1.526 (Iota variant), C.37 (Lambda variant), P.3 (Theta variant), N.10, B.1.1.263 and B.1.1.529 (Omicron variant). Our main findings are discussed further below.

### 4.1 The galectin-like binding pocket in BCoV and SARS-CoV-2 is evolving: a consequence of novel insertions and loops

The NTD of SARS-CoV-2 shares a similar fold to the human Galectin-3 domain (CATH topology (2.60.120), which is known to bind sugars. In this study, we predicted four binding pockets (pockets 1-4) in NTD of SARS-CoV-2, three of which were previously known. The predicted sugar binding ‘pocket 1’ in SARS-CoV-2, corresponds to the galectin-binding pocket in human Galectin-3. Fantini et al. (2020) identified critical sialic acid-binding residues in this pocket in SARS-CoV-2. Our analyses, comparing SARS-CoV-2 to related coronaviruses, indicates that insertions in NTD of SARS-CoV-2 gave additional loops that extended pocket 1, enhancing contact and binding with sialic acid. Other indel regions that emerged in SARS-CoV-2 NTD co-localise at or near the other putative sugar binding pockets, possibly also tuning the interactions of these pockets with sialic acids.

### 4.2 Pockets 2 and 3 are diversity hotspots and potential targets for the ongoing evolution of the virus and increased infectivity

Pockets 2 and 3 were found to be even more structurally variable, across the BCoVs than pocket 1. These pockets have been shown by our studies and others (Awasthi et al., 2020) to be evolutionary hotspots due to significant indel innovations. Our work highlights how the IR2 indel region can affect both pockets 2 and 3 simultaneously since the loop sits between these 2 pockets. IR2 contains a well-known sugar binding motif for sialic acids (72-GTNGTKR-78 motif) that is conserved in different BCoVs supporting a functional role for this region in infection (Behloul et al., 2020). Awasthi et al. (2020), also showed by docking and simulations using sialiosides and NTD structures of SARS-CoV-2 and MERS-CoV that the flexibility of the loop (formed by 244L-261G in pocket 2) could enable binding to a wide range of sialosides.

Recent experimental studies have suggested that the deletions (at positions 69 and 70) in pocket 3 in the SARS-CoV-2 Alpha variant affect the infectivity of the virus (Kemp et al., 2021; Liu et al., 2021). Important to note is that these positions (69-70) bridge both pockets 2 and 3. Notably, a recent study in Brazil (Resende et al., 2021) reported communitary transmission of different lineages of the variant of concern (VOC) gamma (P.1), which harbours NTD indels 69-70 in pocket 3 and an insertion at 214, also in pocket 3. These VOCs were thought to be responsible for the widespread transmission of SARS-CoV-2 in Brazil. The same study also reported a new variant of interest (VOI) N.10, emerging from the B.1.1.33, which harbours NTD deletions that map to both pocket 2 (residues 256-258) and pocket 3 (residue 211) and suggests that these newly emerging variants are likely to be more resistant to antibody neutralisation than the parental variants of concern. An even more recent VOC, Omicron, also has mutations and deletions in pocket 2 and 3 suggesting similar issues and reasons for concern.

All of the structural innovations that change the binding pockets may have an important role in altering sialic acid binding thereby changing the infectivity profile. Recent SARS-CoV-2 variants show wide changes in infectivity (Dicken et al., 2021). The potential role of the NTD in altered infectivity is supported by our binding energy studies which predicted that some VOC variants, particularly in pocket 3, could increase binding to sugars, discussed again in detail below.

Some of these changes could represent an ongoing adaptation of the virus to human cells with their specific sugar modifications. These observations combined with the Dicken et al. study (2021) on increased efficiency of cell entry for the Alpha VOC supports the need for further studies to characterise these binding pockets and monitor any ongoing (pocket 2 and 3) changes as new VOCs emerge.

### 4.3 Differences in sugar-binding affinities across the coronaviruses, and in the variants of concern/interest in SARS-CoV-2

We used established computational tools (HADDOCK and its scoring tool PRODIGY) to analyse changes in binding energy for different BCoVs binding to sialic acid. We found that the NTD domain in the Spike protein of SARS-CoV-2 binds sialic acid more strongly than SARS-CoV, in sugar-binding pockets 1 to 3. Furthermore, some VOCs and VOIs also bind more strongly in pocket 1 (Kappa variant; E154K) and pocket 3 (T95I found in Delta, Iota, Mu variants), compared to the original Wuhan-1 strain of SARS-CoV-2 and SARS-CoV (see Table 5). T95I mutation has been reported to be one of the most prevalent mutations in the new (Delta Plus) strains of the Delta lineage (namely AY.2 or B.1.617.2.1) (Kannan et al., 2021). It is worth noting that the proximity of mutations to binding pockets may also alter the accessibility to the binding pocket or may alter binding to other regions proximal to the pocket (sialic acids are found as part of bigger macromolecules) even if not directly altering sialic acid binding energy.

We propose continuous monitoring of NTD mutations and indels in the context of newly emerging variants and their impacts on sugar-binding. For example, as regards pocket 1 and 3, our studies found that newly emerging strains such as from Brazil (P.1), South Africa (B.1.1.263 and Omicron B.1.1.529) and Delta plus (AY.2) have enhanced binding energy (as compared to original Wuhan strain) that may be associated with higher levels of transmission. Our structural study helps to rationalise potential effects of new VOCs and provide hypotheses for experiments. For example, the Omicron variant has a rather unique 3 amino acid insertion at position 214 which lies near to pocket-3.

Concern regarding the putative increase in transmission of the Omicron VOC highlights the need for further experimental characterisation of the roles of these NTD pockets in infection. In particular, the dramatic changes in virus entry efficiency achieved by changes to IR2 should be studied further, especially by experimental study and in relation to pockets 2 & 3, to help rationalise likely impacts of emerging VOCs.

### 4.4 Predicted pockets: implication in drug design

We observe high conservation in pocket 1 (galectin-like binding pocket, structural similarity score of 83.50 out of 100) and pocket 4 (score 93.80 out of 100). The druggable nature of pocket 1 is also supported by the study by Fantini et al. (2020) which showed that the drugs chloroquine and hydroxychloroquine fully mimic the way in which the sialic acid binds in pocket 1 of NTD, and in the presence of these drugs SARS-CoV-2 no longer binds to sialic acid. Dually targeting galectins and the sialic acid-binding domain of SARS-CoV-2 has been suggested as a promising strategy for COVID-19 for preventing viral entry and modulating the host immune response (Caniglia et al., 2020).

In addition to pocket 1, we suggest targeting pocket 4, which is much more highly conserved among coronaviruses. Designing inhibitors that target structurally conserved regions or pockets in the BCoVs, such as the conserved sialic acid binding pockets (1 and 4) found in SARS-CoV-2, seems to be a promising strategy to inhibit protein function and block viral entry (Caniglia et al., 2020).

## 5 Conflict of Interest

The authors declare that the research was conducted in the absence of any commercial or financial relationships that could be construed as a potential conflict of interest.

## 6 Author Contributions

C.O., J. L. conceived the idea. S.D.L., C.O., J.L. designed the experiments. S.D.L. and V.P.W. performed the experiments. All authors analysed and interpreted the results, wrote this manuscript, performed the revisions.

## 7 Funding

S.D.L. is funded by a Fundamental Research Grant Scheme from the Ministry of Higher Education Malaysia [FRGS/1/2020/STG01/UKM/02/3]. V.P.W. is funded by Biotechnology and Biological Sciences Research Council (BBSRC) [BB/W003368/1]. The APC was funded by BBSRC.

## 8 Supplementary Material

The Supplementary Material for this article can be found online at:

## 9 Data Availability Statement

The original contributions presented in the study are included in the article/supplementary material, further inquiries can be directed to the corresponding author/s.

## Notes

### Competing Interest Statement

The authors have declared no competing interest.

