## Supplementary File for "Insertions in the SARS-CoV-2 Spike N-Terminal Domain May Aid COVID-19 Transmission"

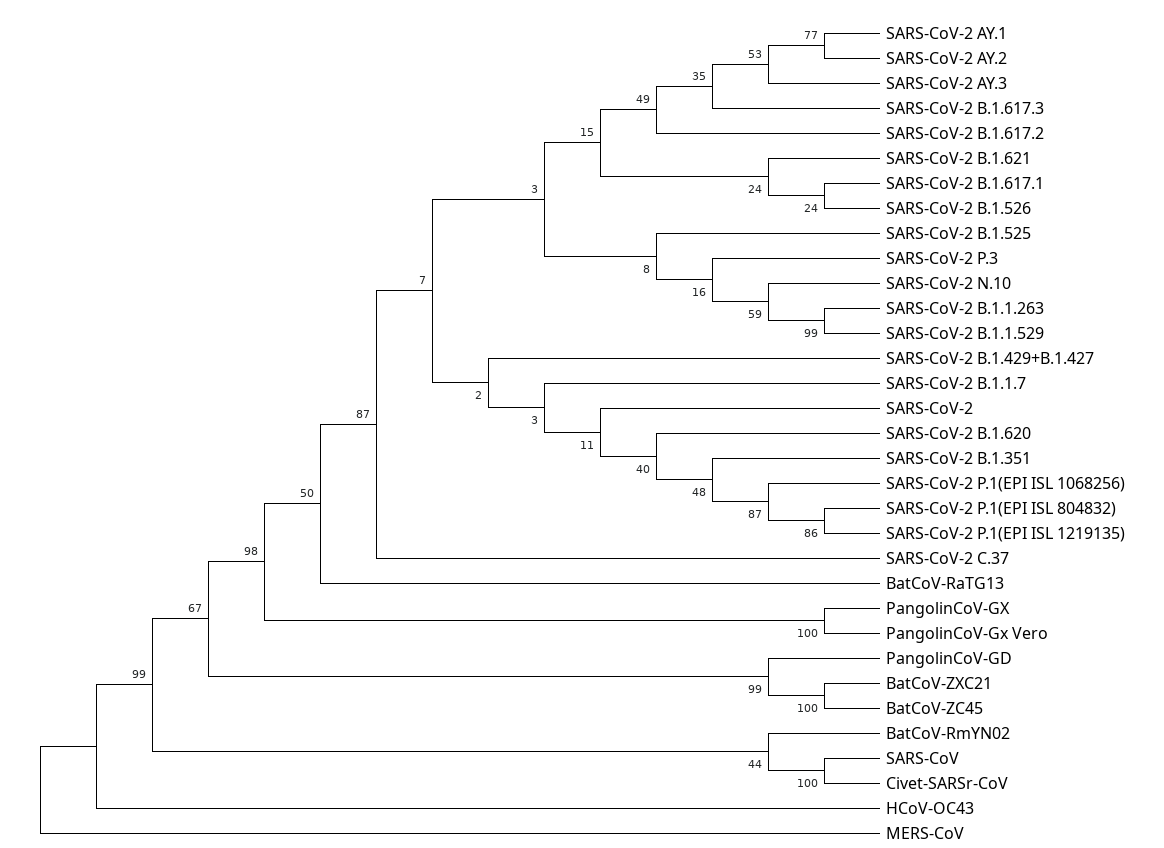

**Supplementary Figure 1.** Bootstrap consensus tree of BCoV NTD amino acid sequences. The phylogenetic tree was inferred according to the Maximum Likelihood method. Genetic distance was computed using the Whelan And Goldman model and gamma-distributed rate variation among sites (WAG + G). The bootstrap consensus tree inferred from 1000 replicates.

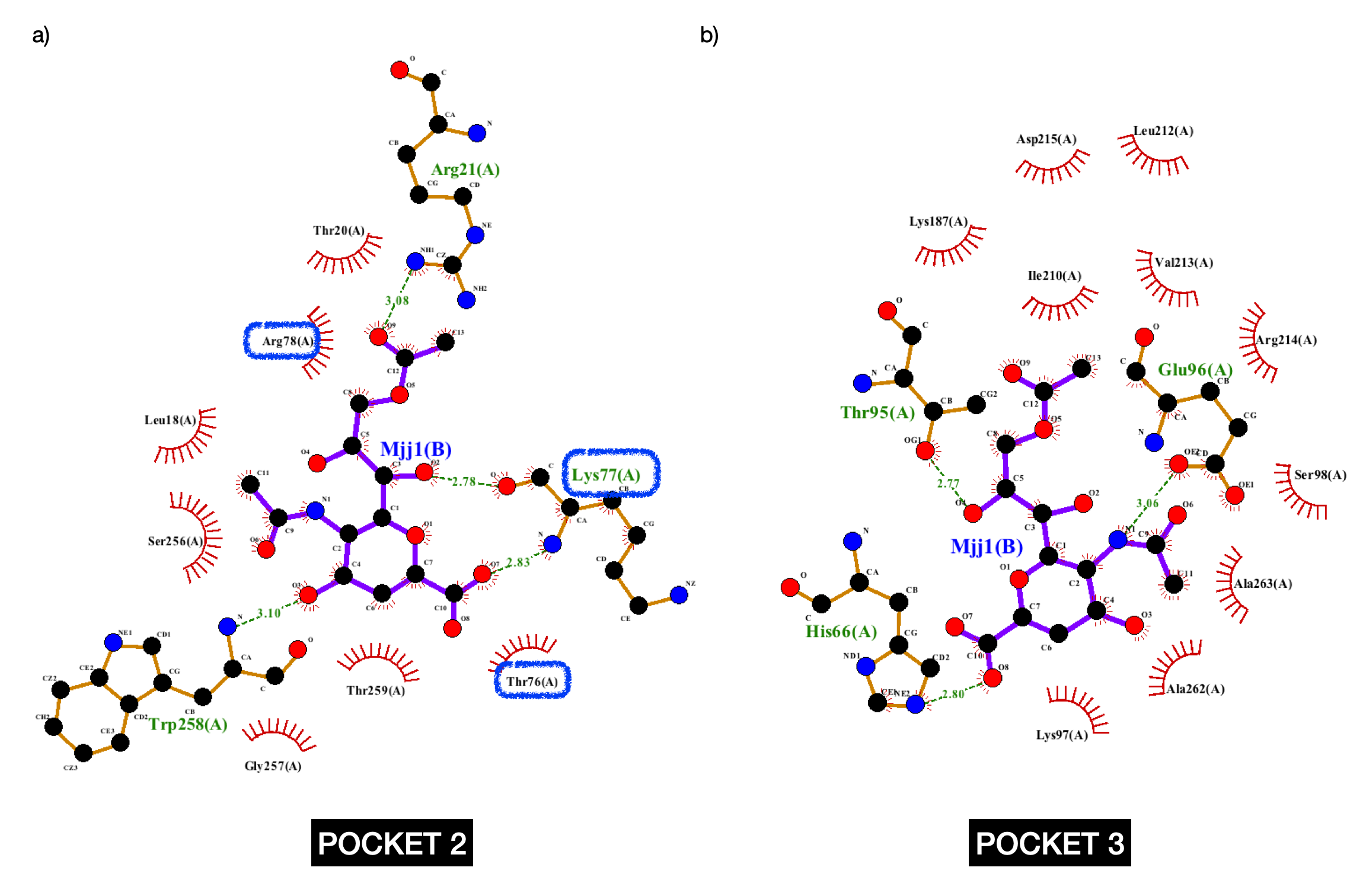

**Supplementary Figure 2.** LigPlot showing H-bonding interactions of sialic acid with pocket 2 and pocket 3 of SARS-CoV2 NTD. Residues highlighted in blue involve the sugar binding motif (72-GTNGTKR-78).

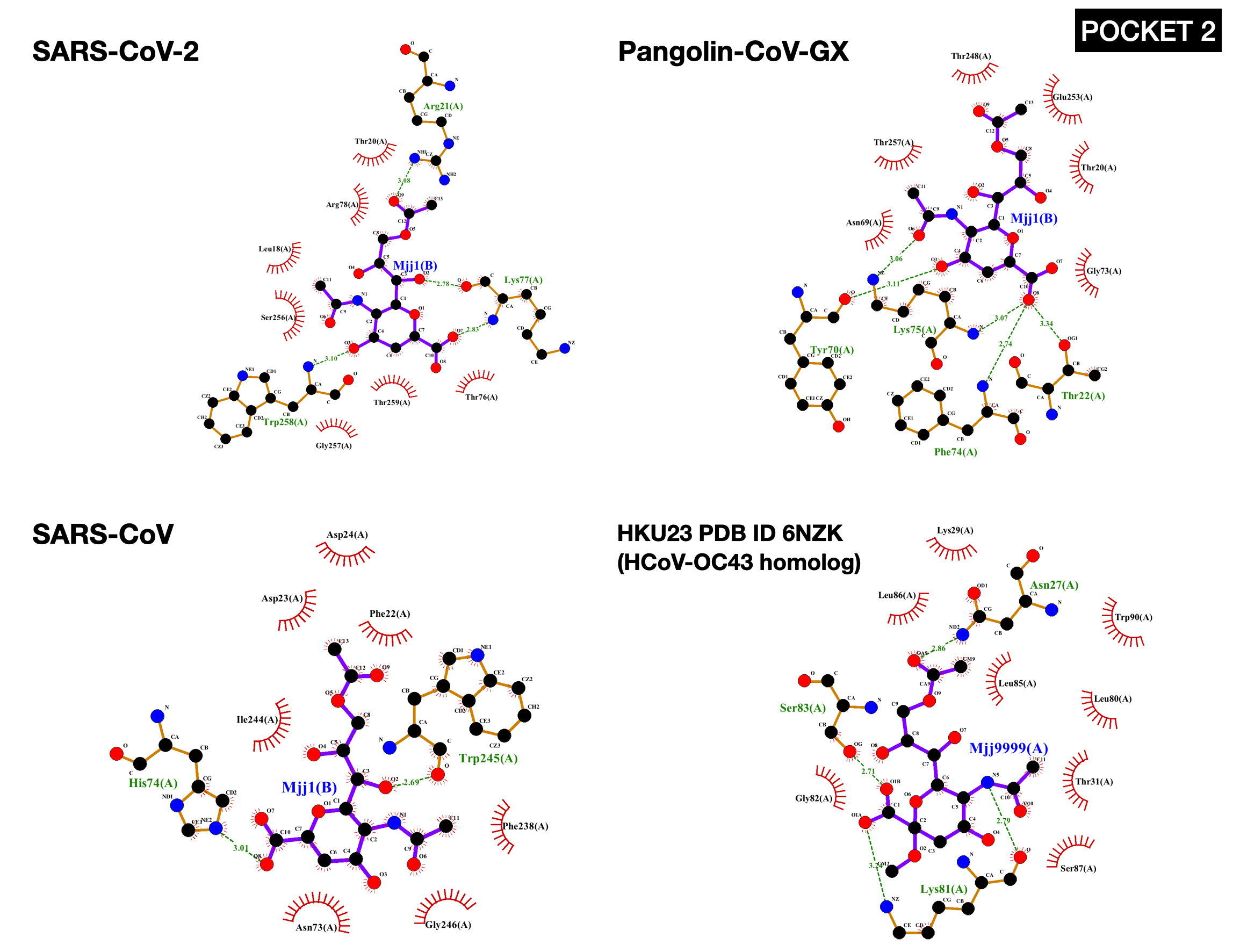

**Supplementary Figure 3.** LigPlots of SARS-CoV-2 NTD, Pangolin-Cov_GX NTD, SARS-CoV NTD and HCoV-OC43 homolog with sialic acid (Pocket 2).

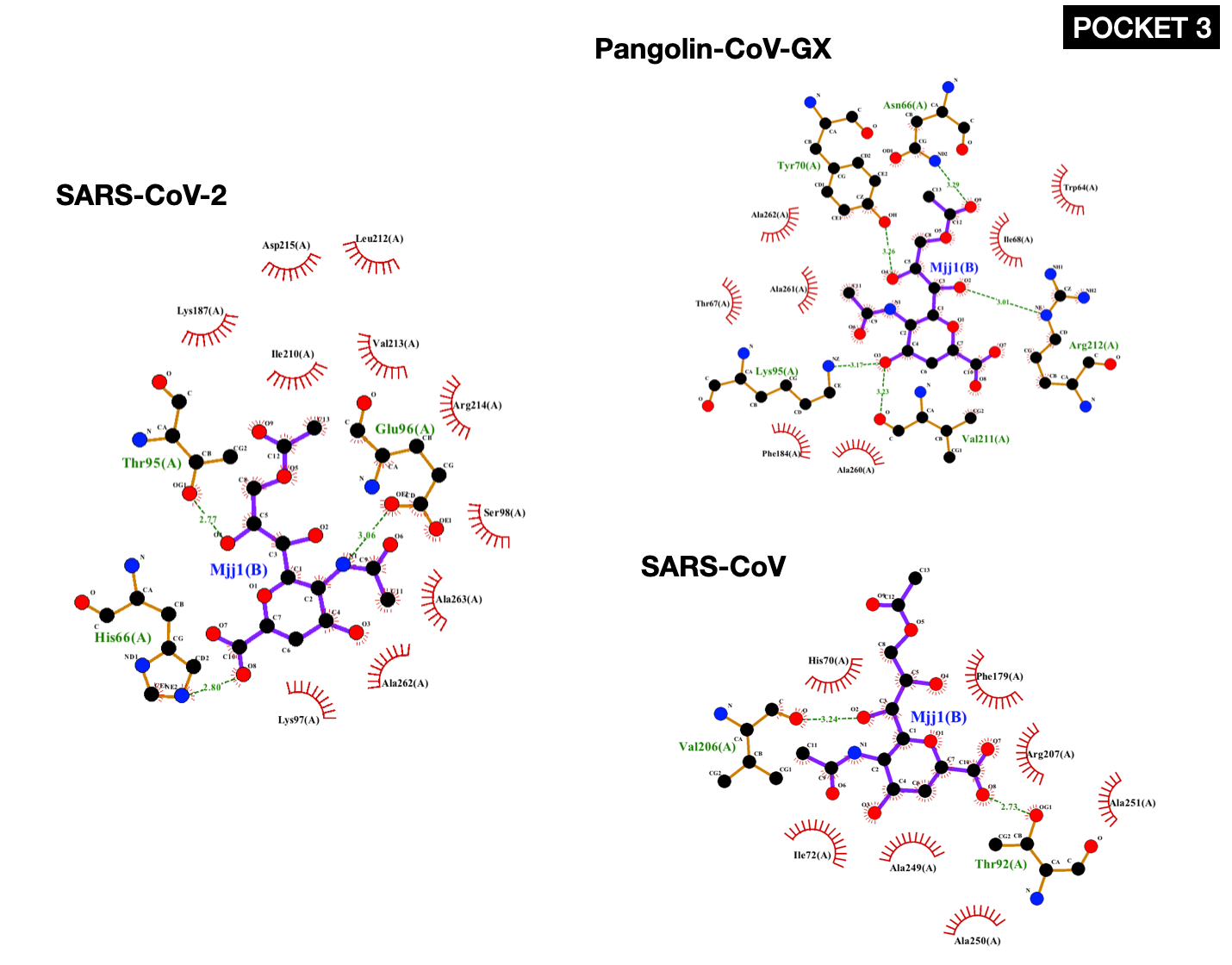
 **Supplementary Figure 4.** LigPlots of SARS-CoV-2 NTD, Pangolin-Cov_GX NTD and SARS-CoV NTD with sialic acid (Pocket 3).

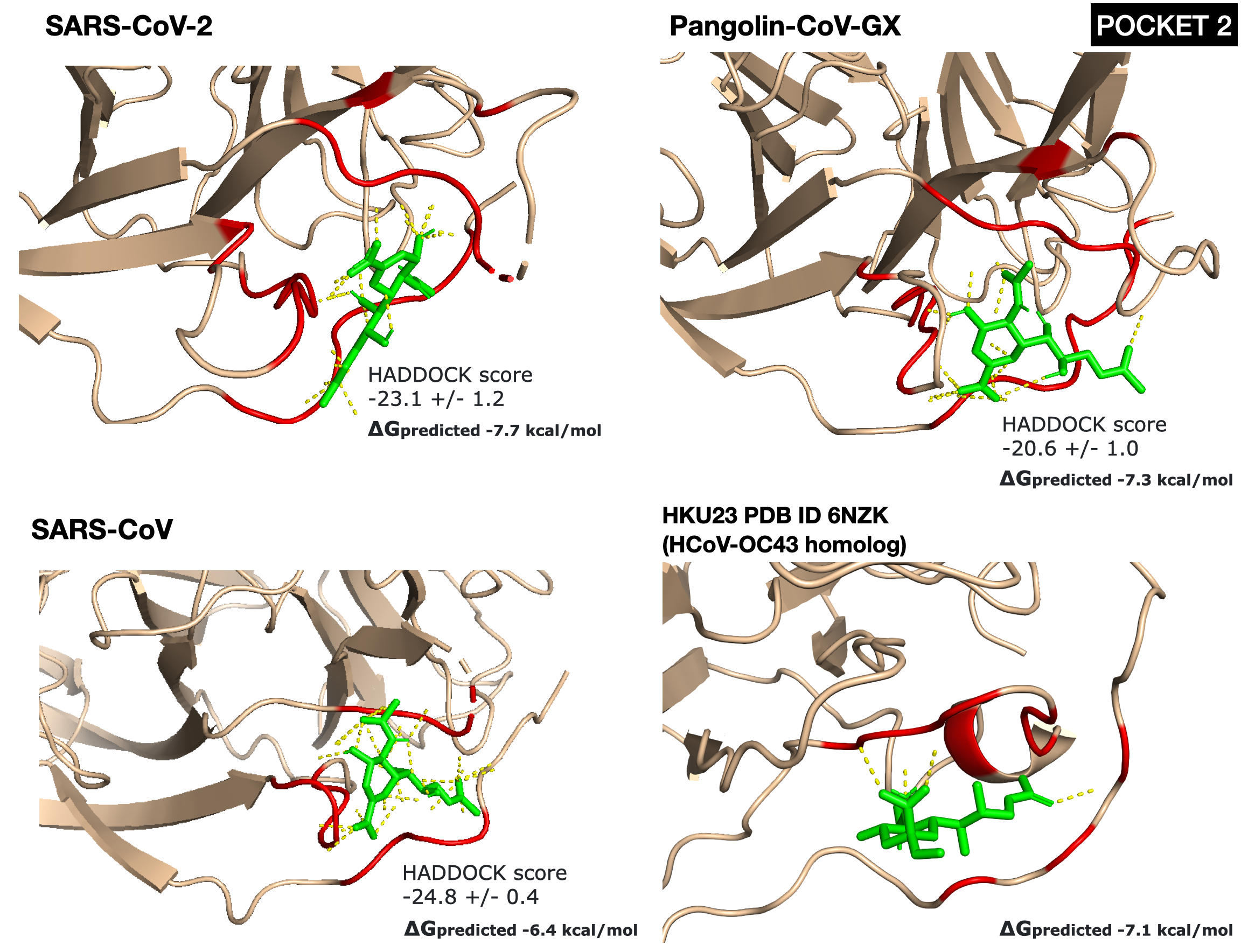

**Supplementary Figure 5.** HADDOCK sialic acid docking result of SARS-CoV-2 NTD, Pangolin-CoV-GX NTD, SARS-CoV NTD and experimental NTD structure of HCoV-OC43 homolog with sialic acid bound. The HADDOCK score is shown together with the PRODIGY predicted binding affinity. The sugar binding residues are shown in red.

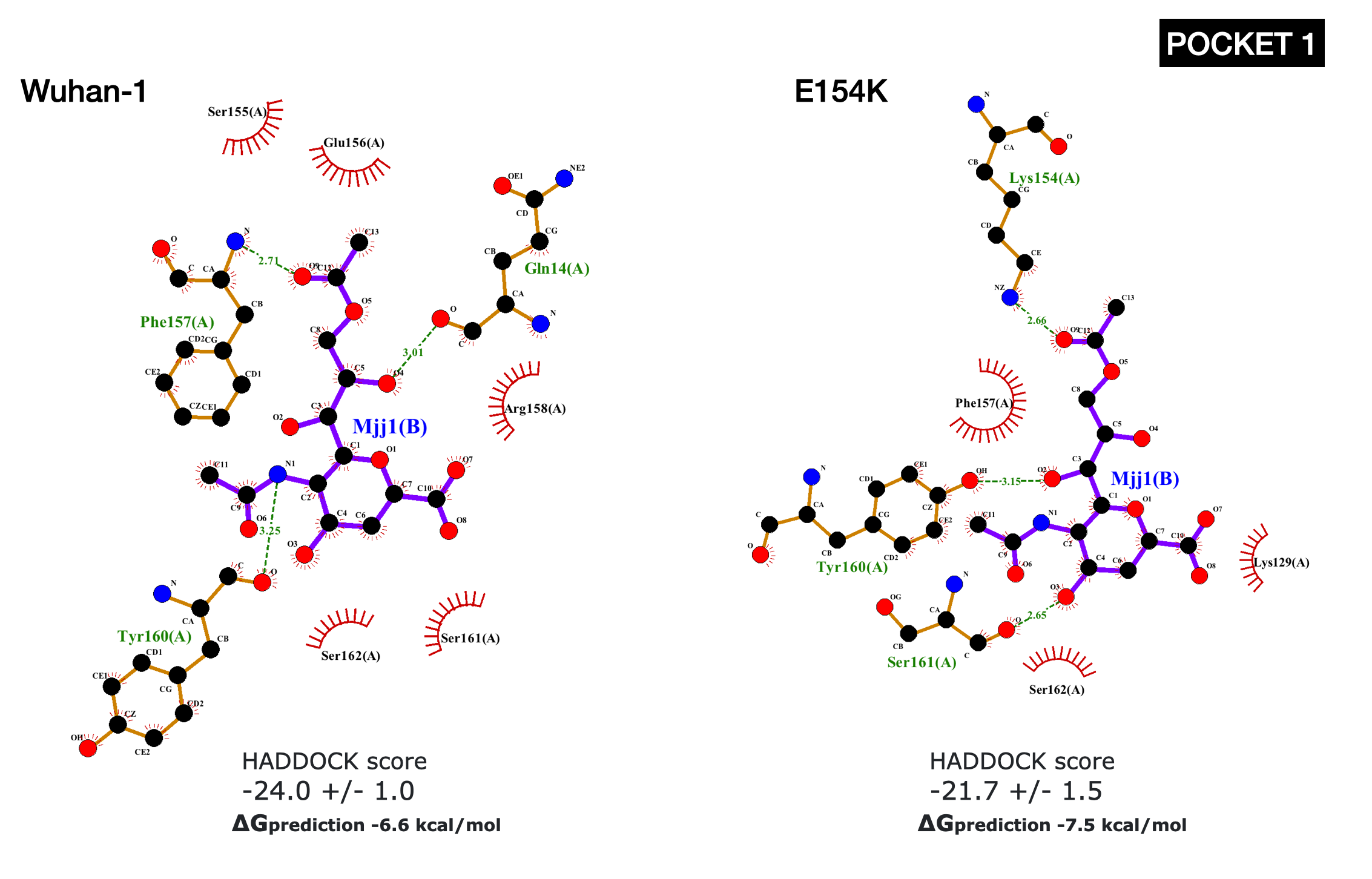

**Supplementary Figure 6.** LigPlots of SARS-COV-2 Wuhan-1 NTD and SARS-COV-2 E154 NTD mutant with sialic acid (Pocket 1).

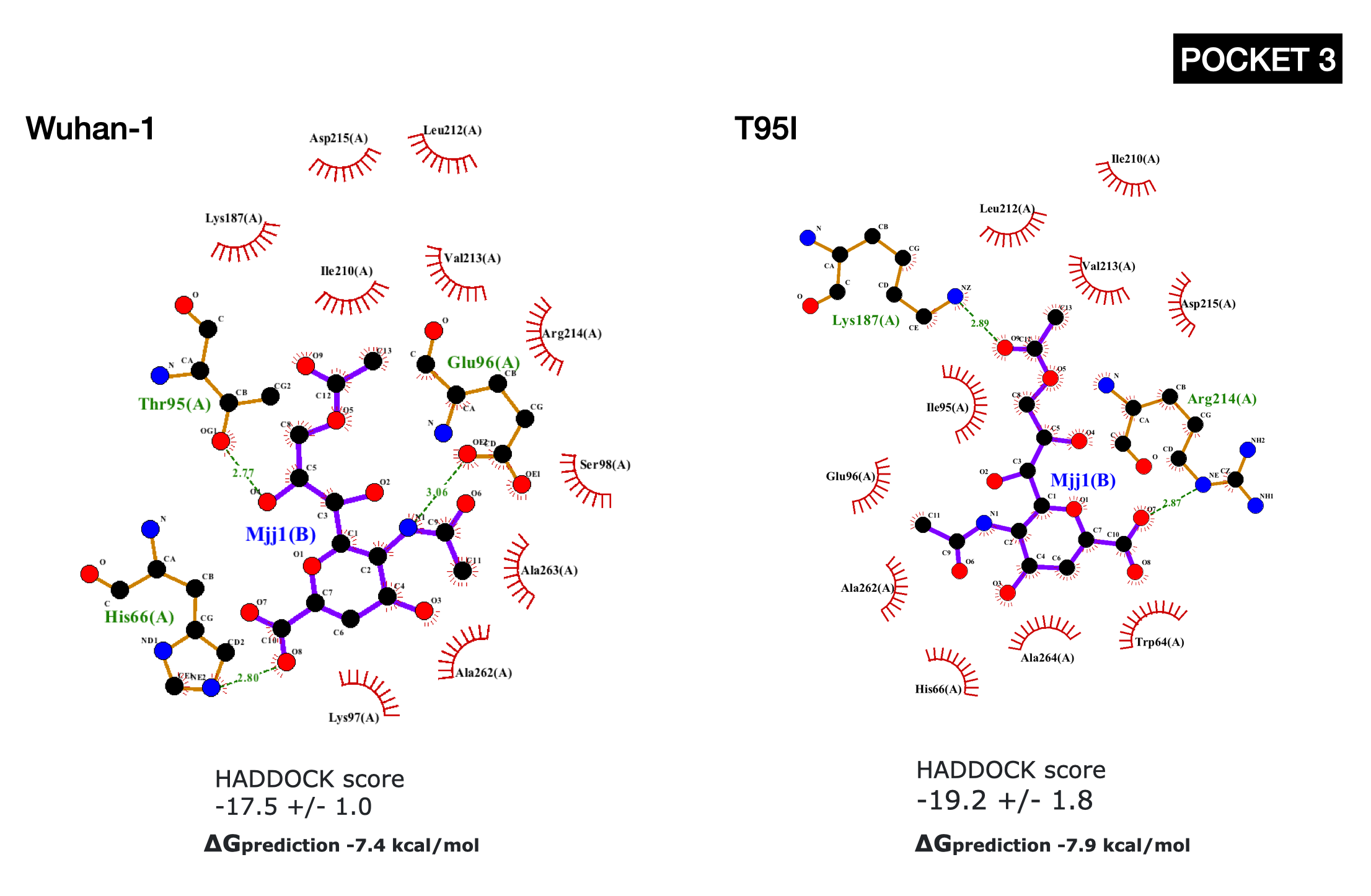

**Supplementary Figure 7.** LigPlots of SARS-COV-2 Wuhan-1 NTD and SARS-COV-2 Wuhan-1 T95I NTD mutant with sialic acid (Pocket 3).

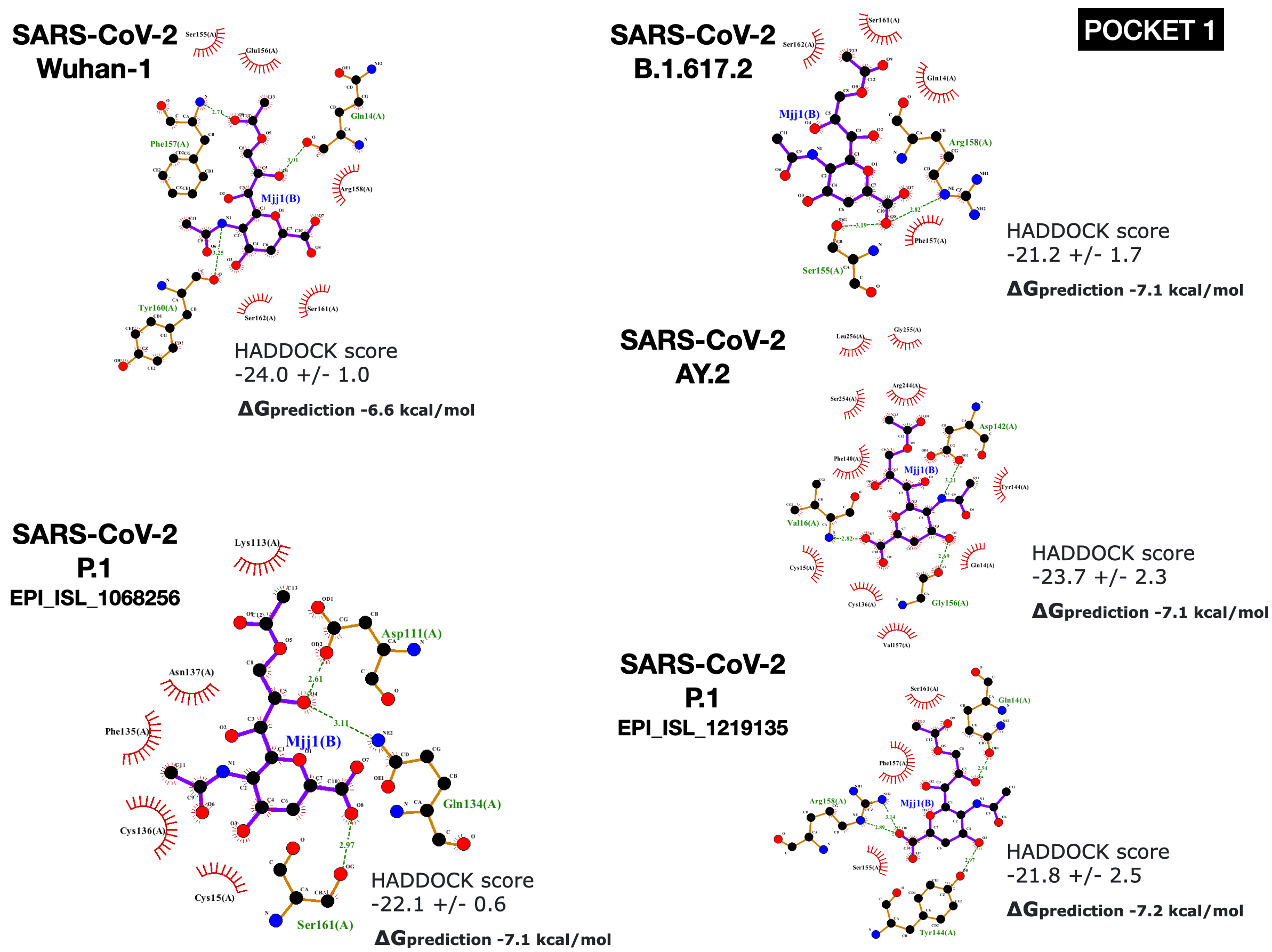

**Supplementary Figure 8.** LigPlots of SARS-COV-2 Wuhan-1 NTD, SARS-COV-2 B.1.617.2 NTD, SARS-CoV-2 AY.2 and SARS-COV-2 P.1 (EPI_ISL_1068256 and EPI_ISL_1068256) with sialic acid (Pocket 1).

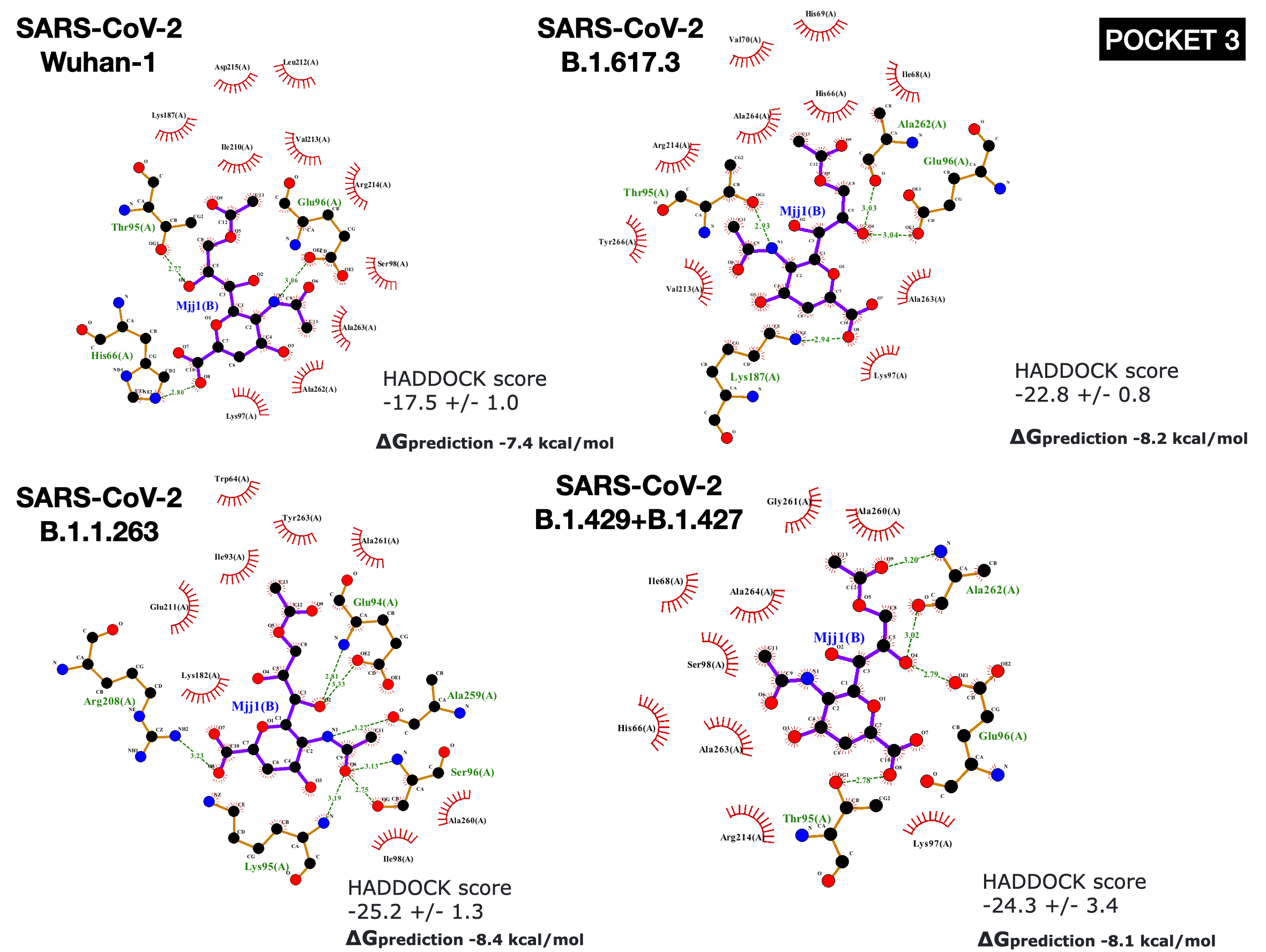

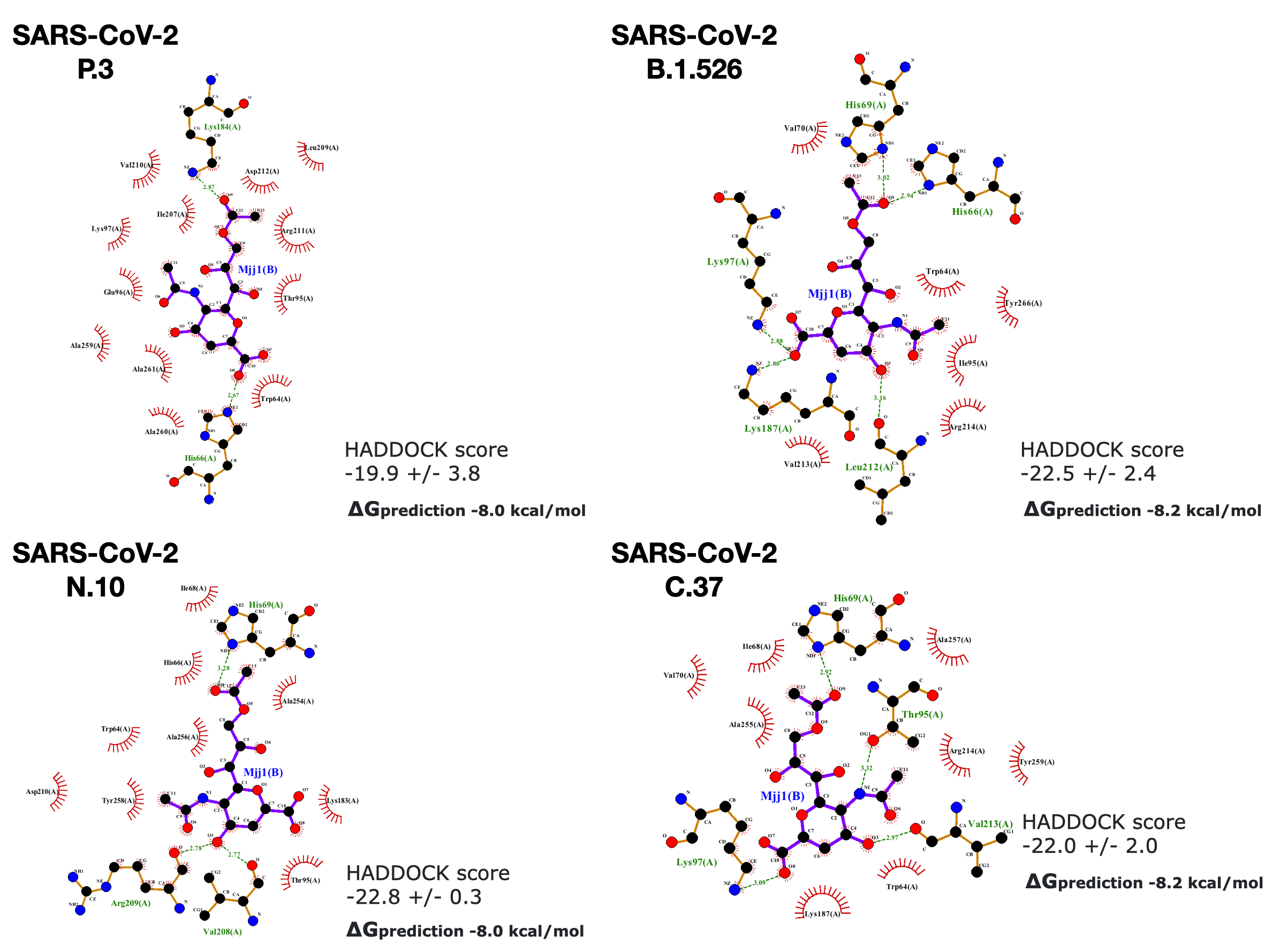

**Supplementary Figure 9.** LigPlots of SARS-CoV-2 Wuhan-1 NTD, SARS-CoV-2 B.1.617.3 NTD, SARS-CoV-2 B.1.429+B.1.427 NTD, SARS-CoV-2 B.1.526 NTD, , SARS-CoV-2 P.3 NTD, SARS-CoV-2 C.37 NTD, SARS-CoV-2 N.10 NTD and SARS-CoV-2 B.1.1.263 with sialic acid (Pocket 3).

**Supplementary Table 1.** NCBI/GISAID ID and common name of coronavirus sequence used in this study.

| **NCBI/GISAID ID** | **Name** |
| --- | --- |
| EPI_ISL_412977 | Bat coronavirus RmYN02 (BatCoV-RmYN02) |
| AVP78042.1 | Bat coronavirus ZXC21 (BatCoV-ZXC21) |
| MG772933.1 | Bat coronavirus ZC45 (BatCoV-ZC45) |
| QHR63300.2 | Bat coronavirus RaTG13(BatCoV-RaTG13) |
| AAU04646.1 | Civet severe acute respiratory syndrome (SARS)-associated coronavirus (Civet-SARSr-Cov) |
| AAP13441.1 | SARS-associated coronavirus (SARS-CoV) |
| QIA48623.1 | Pangolin coronavirus GX (PangolinCoV-GX) |
| QIQ54048.1 | Vero E6 cell passaged Pangolin coronavirus GX (PangolinCoV-GX Vero) |
| QIG55945.1 | Pangolin coronavirus GD (PangolinCoV-GD) |
| AAR01015.1 | Human coronavirus OC43 (HCoV-OC43) |
| KJ156866.1 | Middle East respiratory syndrome–related coronavirus (MERS-CoV) |
| YP_009724390.1 | SARS-CoV-2 (Wuhan-Hu-1 strain) |
| EPI_ISL_741243 | SARS-CoV-2 B.1.1.7 (Alpha variant) |
| EPI_ISL_736967 | SARS-CoV-2 B.1.351 (Beta variant) |
| EPI_ISL_804832 | SARS-CoV-2 P.1 (Gamma variant) |
| EPI_ISL_1068256 | SARS-CoV-2 P.1 (Gamma variant) |
| EPI_ISL_1219135 | SARS-CoV-2 P.1 (Gamma variant) |
| EPI_ISL_2521822 | SARS-CoV-2 B.1.617.1 (Kappa variant) |
| EPI_ISL_2521781 | SARS-CoV-2 B.1.617.2 (Delta variant) |
| EPI_ISL_6656070 | SARS-CoV-2 AY.1 (Delta Plus variant) |
| EPI_ISL_5811040 | SARS-CoV-2 AY.2 (Delta Plus variant) |
| EPI_ISL_6695006 | SARS-CoV-2 AY.3 (Delta Plus variant) |
| EPI_ISL_1970532 | SARS-CoV-2 B.1.617.3 |
| EPI_ISL_2801732 | SARS-CoV-2 B.1.429+B.1.427 (Epsilon variant) |
| EPI_ISL_3692979 | SARS-CoV-2 B.1.526 (Iota variant) |
| EPI_ISL_3574438 | SARS-CoV-2 B.1.525 (Eta variant) |
| EPI_ISL_3147606 | SARS-CoV-2 B.1.620 |
| EPI_ISL_3654107 | SARS-CoV-2 B.1.621 (Mu variant) |
| EPI_ISL_2535683 | SARS-CoV-2 P.3 (Theta variant) |
| EPI_ISL_3672611 | SARS-CoV-2 C.37 (Lambda variant) |
| EPI_ISL_1181371 | SARS-CoV-2 N.10 |
| EPI_ISL_6640916 | SARS-CoV-2 B.1.1.263 |
| EPI_ISL_6704875 | SASR-CoV-2 B.1.1.529 (Omicron variant) |

**Supplementary Table 2.** Structure modelling template, structural template sequence identity and model quality (normalised DOPE) of BCoV Spike protein NTD models using FunMod modelling platform

| **NTD model** | **Template** | **Sequence Identity** | **Normalised DOPE** |
| --- | --- | --- | --- |
| BatCoV-RmYN02 | 7CN8 | 47% | -0.36 |
| BatCoV-ZXC21 | 7CN8 | 65% | -0.86 |
| BatCoV-ZC45 | 7CN8 | 63% | -0.80 |
| Civet-SARSr-Cov | 6ACC | 100% | -0.60 |
| Pangolin-CoV-Gx-Vero | 7CN8 | 97% | -0.58 |
| SARS-CoV-2 B.1.1.7 | 7C2L | 98 % | -0.39 |
| SARS-CoV-2 B.1.351 | 7C2L | 95 % | -0.44 |
| SARS-CoV-2 P.1 (EPI_ISL_804832) | 7C2L | 99 % | -0.45 |
| SARS-CoV-2 P.1 (EPI_ISL_1068256) | 7C2L | 98 % | -0.37 |
| SARS-CoV-2 P.1 (EPI_ISL_1219135) | 7C2L | 97 % | -0.31 |
| SARS-CoV-2 B.1.617.1 | 7C2L | 99 % | -0.37 |
| SARS-CoV-2 B.1.617.2 | 7C2L | 99 % | -0.37 |
| SARS-CoV-2 AY.1 | 7C2L | 96 % | -0.31 |
| SARS-CoV-2 AY.2 | 7C2L | 97% | -0.31 |
| SARS-CoV-2 AY.3 | 7C2L | 97 % | -0.32 |
| SARS-CoV-2 B.1.617.3 | 7C2L | 99 % | -0.43 |
| SARS-CoV-2 B.1.429+B.1.427 | 7C2L | 99 % | -0.39 |
| SARS-CoV-2 B.1.526 | 7C2L | 99 % | -0.34 |
| SARS-CoV-2 B.1.525 | 7C2L | 97 % | -0.47 |
| SARS-CoV-2 B.1.620 | 7C2L | 94 % | -0.51 |
| SARS-CoV-2 B.1.621 | 7C2L | 83 % | -0.19 |
| SARS-CoV-2 P.3 | 7C2L | 98 % | -0.33 |
| SARS-CoV-2 C.37 | 7C2L | 99 % | -0.44 |
| SARS-CoV-2 N.10 | 7C2L | 96% | -0.46 |
| SARS-CoV-2 B.1.1.263 and B.1.1.529 | 7C2L | 93% | -0.48 |

**Supplementary Table 3.** Predicted lDDT scores of AlphaFold2 BCoV Spike protein NTD models. Models with predicted IDDT score above 70 are considered to be acceptable.

| **UniProt ID** | **lDDT score** |
| --- | --- |
| A0A088DJY6 | 79.65 |
| A0A0A7UZR7 | 95.36 |
| A0A0K1Z074 | 94.93 |
| A0A0K2RVL1 | 93.97 |
| A0A0U1UYX4 | 94.32 |
| A0A0U1WJY8 | 89.79 |
| B7U2N3 | 93.06 |
| B8Q8S6 | 85.63 |
| C0KZ00 | 89.85 |
| E0XIZ3 | 94.14 |
| H9AA65 | 94.41 |
| H9BZX9 | 88.48 |
| P11224 | 95.65 |
| P36334 | 95.64 |
| Q06BD7 | 94.19 |
| Q0Q475 | 93.45 |
| Q19U46 | 89.49 |
| Q77NQ7 | 91.86 |
| Q8BB25 | 96.78 |
| Q8JSP8 | 96.58 |
| R9QTA0 | 88.90 |

**Supplementary Table 4**. Residues in the known and putative sugar binding pockets of SARS-CoV-2

| **Function** | **Note** | **Viruses that contain the pocket** | **Pocket** | **Pocket Residues** | **Citations** |
| --- | --- | --- | --- | --- | --- |
| Sialic acid/ ganglioside binding | Sialic acid-binding region (*In silico* structural and molecular modelling study) | Identified by analysis of SARS-CoV-2 | 1 | D111, S112, K113, Q134, F135, C136, N137, F140, G142, E156, F157, R158, Y160, S161, S162 | (Fantini et al., 2020) |
| Sugar binding | BcoV sugar-binding domain (identified by analysis of bovine coronavirus) | BovineCoV-NTD, PHEV-NTD, HCoV-OC43-NTD, HCoV-HKU23-NTD, HKU1-NTD, MHV-NTD | 1 | E154, F157, Y160 | (Cheng et al., 2019; Behloul et al., 2020) |
| Sugar binding | Sugar receptor-interacting motif (identified by analysis of bacteriophage CBA120, infectious bronchitis coronavirus) | BovineCoV-NTD, PHEV-NTD, HCoV-OC43-NTD, HCoV-HKU23-NTD, HKU1-NTD, MHV-NTD | 2, 3 | G72, T73, N74, G75, T76, K77, R78 | (Cheng et al., 2019; Behloul et al., 2020) |
| Sialic acid-binding | Identified by analysis of SARS-CoV-2 – similar to those reported for HCoV-OC43 structure with sialic-acid bound | BovineCoV-NTD, PHEV-NTD, HCoV-OC43-NTD, HCoV-HKU23-NTD, HKU1-NTD, MHV-NTD | 2 | R21, Q23, L24, H69, F79, P82, R246 | (Cheng et al., 2019; Tortorici et al., 2019; Baker et al., 2021) |
| Druggable Pocket | Predicted using SiteMap (PDB ID 7JJI) | Identified by analysis of SARS-CoV-2 | 2 | R21, T22, Q23, L24, P26, R78, P82, V83, L110, F135, C136, N137, R237 | (Di Gaetano et al., 2021) |
| Druggable pocket | Predicted using CavityPlus (PDB ID 7C2L) | Identified by analysis of SARS-CoV-2 | 2 | V16, N17, L18, T19, T20, R21, T22, I68, H69, N74, G75, T76, K77, R78, F79, D80, L244, S247, S256, G257, W258, T259, A260 | This study |
| Druggable Pocket | Predicted using SiteMap (PDB ID 7JJI) | Identified by analysis of SARS-CoV-2 | 3 | F92, S94, E96, K97, S98, R102, N121, V126, I128, M177, D178, K182, N188, R190, F192, I203, L226, V227, L229 | (Di Gaetano et al., 2021) |
| Druggable pocket | Predicted using CavityPlus (PDB ID 7C2L) | Identified by analysis of SARS-CoV-2 | 3 | A27, W64, F65, H66, A67, I68, H69, V70, S71, G72, A93, S94, T95, E96, K97, S98, N99, I100, N185, F186, K187, N188, L189, I210, N211, L212, V213, D214, D215, L216, P217, A260, G261, A262, A263, A264, Y265, Y266 | This study |
| Druggable pocket | Predicted using CavityPlus (PDB ID 7C2L) | Identified by analysis of SARS-CoV-2 | 4 | Y38, P39, D40, K41, V42, F43, R44, S45, S46, V47, L48, H49, S50, T51, Q52, D53, T274, F275, L276, L277, K278, Y279, N280, E281, C291, E298, T299, K300, C301, T302, L303 | This study |

**Supplementary Table 5.** NTD mutations found in SARS-CoV-2 Variants of Concern (VOC)/Interest (VOI) and present in NTD sugar binding pockets 1 to 3.

| Name | Variant Classification | NTD mutation/deletion | NTD mutation lies close to sugar-binding pocket |
| --- | --- | --- | --- |
| Alpha variant (B.1.1.7) | VOC | H69del, V70del, Y144del | Pocket 1: Y144del  Pocket 2: H69del, V70del  Pocket 3: H69del, V70del |
| Beta variant (B.1.351) | VOC | L18F, D80A, D215G, L241del, A242del, L243del | Pocket 1: L18F  Pocket 2: L18F, D80A, L241del, L242del, A243del,  Pocket 3: D215G |
| Gamma variant (P.1)  EPI_ISL_804832 | VOC | L18F, T20N, P26S, D138Y, R190S | Pocket 1: L18F, D138Y  Pocket 2: L18F, T20N, D80A  Pocket 3: P26S, R190S |
| Gamma variant (P.1)  EPI_ISL_1219135 | VOC | L18F, T20N, P26S, D138Y, N188S, L189del, R190del | Pocket 1: L18F, D138Y  Pocket 2: L18F, T20N,  Pocket 3: P26S, N188S, L189del, R190del |
| Gamma variant (P.1)  EPI_ISL_1068256 | VOC | L18F, P26S, D138Y, ins214ANRN | Pocket 1: L18F, D138Y  Pocket 2: L18F  Pocket 3: P26S, ins214ANRN |
| Kappa variant  (B.1.617.1) | VOI | T95I, E154K | Pocket 1: E154K  Pocket 3: T95I |
| Delta variant (B.1.617.2) | VOC | T19R, T95I | Pocket 2: T19R  Pocket 3: T95I |
| Delta Plus variant  (AY.1) | VOC | T19R, H49Y, T95I, G142D, E156G, F157del, R158del, W258L | Pocket 1: T19R, G142D, E156G, F157del, R158del  Pocket 2: T19R, W258L  Pocket 3: T95I |
| Delta Plus variant  (AY.2) | VOC | T19R, T95I, G142D, E156G, F157del, R158del, W258L | Pocket 1: T19R, E156G, F157del, R158del  Pocket 2: T19R, W258L  Pocket 3: T95I |
| Delta Plus variant  (AY.3) | VOC | T19R, E156G, F157del, R158del | Pocket 1: T19R, E156G, F157del, R158del  Pocket 2: T19R |
| B.1.617.3 | VOI | T19R, G142D | Pocket 1: T19R, G142D  Pocket 2: T19R |
| Epsilon variant (B.1.429+B1.427) | VOI | S13I (signal peptide), W152C | Pocket 1: W152C |
| Eta variant (B.1.525) | VOI | Q52R, A67V, H69del, V70del, Y144del | Pocket 1: Y144del  Pocket 2: A67V, H69del, V70del  Pocket 3: A67V, H69del, V70del |
| Iota variant (B.1.526) | VOI | L5F (signal peptide), T95I, D253G | Pocket 2: D253G  Pocket 3: T95I |
| B.1.620 | VOI | P26S, H69del, V70del, V126A, Y144del, L242del, A243del, L244del, H245Y | Pocket 1: Y144del  Pocket 2: H69del, V70del, L242del, A243del, L244del, H245Y  Pocket 3: H69del, V70del, P26S |
| B.1.621 | VOI | T95I, Y144S, Y145N | Pocket 1: Y144S, Y145N  Pocket 3: T95I |
| Lambda variant (C.37) | VOI | G75V, T76I, R246del, S247del, Y248del, L249del, T250del, P251del, G252del, D253N | Pocket 2: G75V, T76I, R246del, S247del, Y248del, L249del, T250del, P251del, G252del, D253N |
| Theta variant (P.3) | VOI | L141del, G142del, V143del | Pocket 1: L141del, G142del, V143del |
| N.10 |  | P9L (signal peptide), L141del, G142del, V143del, Y144del, I210V, L212I, N211del, S256del, G257del, W258del | Pocket 1: L141del, G142del, V143del, Y144del  Pocket 2: S256del, G257del, W258del  Pocket 3: I210V, L212I, N211del, |
| B.1.1.263 and  Omicron variant (B.1.1.529)  (Both strains have identical NTD mutations) | VOC | A67V, H69del, V70del, T95I, G142D, V143del, Y144del, Y145del, N211del, L212I, ins214EPE | Pocket 1: G142D, V143del, Y144del, Y145del  Pocket 2: H69del, V70del,  Pocket 3: A67V, H69del, V70del, T95I, N211del, L212I, ins214EPE |

**Supplementary Table 6.** PRODIGY predicted binding energy of sialic acid to SARS-CoV-2 individual mutants, NIL – residue not covered by the structure. Red highlights binding energy values for those variants which increase binding above 0.5 kcal/mol.

| Pocket | Wuhan-1 predicted binding energy | Mutations found in Variant of Concern/Interest | Mutant predicted binding energy |
| --- | --- | --- | --- |
| 1 | -6.6 kcal/mol | L18F (Beta, Gamma)  T19R (Delta, AY.1, AY.2, AY.3, B.1.617.3)  D138Y (Gamma)  G142D (AY.1, AY2. B.1.617.3)  Y144S, Y145N (B.1.621)  W152C (Epsilon)  E154K (Kappa)  E156G (AY.1) | -7.0 kcal/mol  -6.5 kcal/mol  -6.8 kcal/mol  -6.9 kcal/mol  -6.5 kcal/mol  -6.7 kcal/mol  -7.5 kcal/mol  -6.6 kcal/mol |
| 2 | -7.7 kcal/mol | L18F (Beta, Gamma)  T19R (Delta, B.1.617.3)  T20N (Beta, Gamma)  A67V (Eta)  G75V, T76I (Lambda)  D80A (Beta)  H245Y (B.1.620)  D253G (Iota)  D253N (Lambda)  W258L (AY.1, AY.2) | -7.8 kcal/mol  -7.4 kcal/mol  -7.8 kcal/mol  -7.4 kcal/mol  -7.4 kcal/mol  -7.6 kcal/mol  -7.6 kcal/mol  NIL  NIL  -7.4 kcal/mol |
| 3 | -7.4 kcal/mol | P26S (Beta, Gamma)  A67V (Eta, B.1.1.263)  T95I (Kappa, Delta, AY.1, AY.2, Iota, Mu, B.1.1.263)  R190S (Beta, Gamma)  D215G (Beta) | -7.4 kcal/mol  -7.4 kcal/mol  -7.9 kcal/mol  -7.6 kcal/mol  -7.4 kcal/mol |
